# Youth-associated protein TIMP2 alters microglial state and function in the context of aging

**DOI:** 10.1101/2025.05.20.655226

**Authors:** Brittany M. Hemmer, Ana Catarina Ferreira, Sarah M. Philippi, Samuele F. Petridis, Annie Phan, Joseph M. Castellano

## Abstract

There is little understanding of how aging serves as the strongest risk factor for several neurogenerative diseases. Specific neural cell types, such as microglia, undergo age-related maladaptive changes, including increased inflammation, impaired debris clearance, and cellular senescence, yet specific mediators that regulate these processes remain unclear. The aged brain is rejuvenated by youth-associated plasma factors, including tissue inhibitor of metalloproteinases 2 (TIMP2), which we have shown acts on the extracellular matrix (ECM) to regulate synaptic plasticity. Given emerging roles for microglia in these processes, we examined the impact of TIMP2 on microglial function. We show that TIMP2 deletion exacerbates microglial phenotypes associated with aging, including transcriptomic changes in cell activation, increased microgliosis, and increased levels of stress and inflammatory proteins measured in the brain extracellular space by *in vivo* microdialysis. Deleting specific cellular pools of TIMP2 *in* vivo increased microglial activation and altered myelin phagocytosis. Treating aged mice with TIMP2 reversed several phenotypes observed in our deletion models, resulting in decreased microglial activation, reduced proportions of proinflammatory microglia, and enhanced phagocytosis of physiological substrates. Our results identify TIMP2 as a key modulator of age-associated microglia dysfunction. Harnessing its activity may mitigate detrimental effects of age-associated insults on microglia function.

## Introduction

Aging is accompanied by progressive functional decline across multiple organ systems, but its impact on the brain is especially devastating given the dysfunction wrought across key cognitive domains and the increased risk it poses for several neurological disorders. These changes reflect cumulative cellular, molecular, and structural perturbations, including decreased generation of newborn neurons, loss of synaptic and myelin integrity, and microgliosis in vulnerable brain regions like the hippocampus. Understanding mechanisms that regulate aging in the hippocampus is thus essential for identifying new strategies to preserve or restore function across lifespan and to limit risk for neurological disease. Interestingly, the process of aging and AD pose similar challenges to the brain’s immune environment, including aberrant inflammation and elevated exposure to lipid-rich debris from dying cells^1^. The role of microglia, the brain’s innate immune cells, in responding to aging-associated damage has emerged as a promising avenue for investigation, galvanized by the identification of many myeloid-enriched genes associated with AD risk in a series of GWAS over the past decade^2^. While specific states adopted by microglia may initially be protective to limit pathology to surrounding brain tissue, chronic challenges and exposure to debris may create maladaptive responses,^3–6^ which manifests as phagocytic ineptitude^7–10^, production of inflammatory cytokines^11,12^, and increased stress responses^6,13^. Thus, there is a need to identify means to rejuvenate the response of these cells to aging-associated challenges to provide a path for novel therapies that limit the impact of aging on neurological disorders.

Modulating the systemic environment has emerged as a promising strategy to counteract age-related decline in cellular function. Exposure to factors present within young blood rejuvenates the aged brain^14,15^, leading to increased synaptic plasticity^16–18^ and dendritic spine integrity^17^, enhanced vascular remodeling^19^, and improved learning and memory performance in aged mice^16–18^. While much of this work has focused on the impact of blood-borne factors directly on neurons or adult neuroblasts in the hippocampus, recent work has suggested that microglia can take up labeled proteins originating in plasma^20,21^, suggesting that they may be responsive to blood-borne factors. Moreover, in response to treatment with plasma or a platelet factor that acts on immune cells in the periphery, microglia appear to exhibit altered phagocytosis^21^ and activation state^18^, respectively. Despite these intriguing findings, the field lacks detailed characterization of how specific youth-associated factors affect microglial function and how such factors can be harnessed to revitalize the aged brain.

We previously identified tissue inhibitor of metalloproteinases 2 (TIMP2) as a youth-associated blood-borne factor that is capable of revitalizing hippocampal function in aged mice^16^. Levels of TIMP2 are elevated in very young human and mouse plasma and rapidly decline into adulthood and old age^16^. Interestingly, a recent study finds that those expressing *TIMP2* variants associated with higher plasma protein levels exhibit higher cognitive performance in a cohort of aged individuals^22^. Aged mice treated systemically with TIMP2 exhibit similar improvements as mice treated with young plasma, including increased hippocampal plasticity and cognitive improvements, likely as a result of direct TIMP2 action on CNS cells given its entry from blood into brain^16^. While TIMP2 has many canonical and non-canonical targets^23,24^, these studies suggest that TIMP2 has myriad roles within the hippocampal microenvironment, and its role in modulating innate immune function is understudied. We recently reported that TIMP2 plays a critical role in the young hippocampus in regulating memory, synaptic integrity, and adult neurogenesis through modulation of ECM homeostasis^25^, processes that have been found to be regulated by microglia^26,27^. Given the modulation by TIMP2 in processes linked to microglia function and the rejuvenating capacity of TIMP2 in aging contexts, we sought to determine how TIMP2 affects basic microglial function and whether systemic supplementation can restore protective microglial activities that are lost with exposure to age-associated insults.

Here we show that loss of TIMP2 expression from a variety of sources (i.e., removal from global, microglial, and neuronal sources) alters microglia by increasing activation, elevating their inflammatory state, and shifting phagocytosis—all processes associated with aging. Restoring the extrinsic pool of TIMP2 in aged mice through systemic TIMP2 treatment reversed these age-associated phenotypes by reducing microgliosis and specific inflammatory microglial subpopulations and by increasing phagocytosis of physiological substrates in the hippocampus. Together our results demonstrate that the youth-associated protein TIMP2 plays a novel role in modulating microglial function by shifting their responses to accommodate detrimental aging pathologies in a protective fashion. This work firmly positions TIMP2 as a relevant therapeutic target to limit aging pathology and highlights the need for further mechanistic investigation into how systemic factors modulate innate immune function in the brain.

## Results

### Early deletion of TIMP2 alters microglial state

TIMP2 is expressed in cells of the periphery, contributing to high levels in the plasma^16^, and also hippocampal neurons within the CNS^25^, with expression in both compartments declining with age^16^. Interestingly, *Timp2* has been shown to be upregulated in subpopulations of microglia in response to aging and AD pathology as part of a panel of genes termed the disease-associated microglia (DAM) profile^28^. To explore its expression within microglia, we noted its enriched transcript abundance within microglia in published transcriptomic atlases, including in *Tabula Muris Senis*^29^, and in the Allen Brain Cell scRNAseq and MERFISH Atlas^30^, in which *Timp2* is expressed at high levels in the brain, particularly in hippocampal microglia (**Supplementary Fig. 1a-c**). To begin to characterize whether microglia express TIMP2 at the protein level and how its loss influences function at early stages, we isolated primary microglia from neonatal forebrain from wild-type (WT) and TIMP2 knock-out (KO) mice (**Fig. 1a**) at an age when global TIMP2 levels are high^16^. We find high levels of TIMP2 protein by both immunoblotting of primary microglia lysates and immunocytochemistry (ICC), whereas TIMP2 was not detected in TIMP2 KO microglia (**Fig. 1b-c**). We next sought to understand what pathways may be disrupted by its loss in microglia by performing bulk RNA-sequencing on WT and KO primary microglia (**Fig. 1a**). As expected, KO microglia lacked *Timp2* transcript (**Fig. 1d**), and we detected a number of differentially expressed genes in KO microglia relative to WT (252 upregulated, 316 downregulated; **Supplementary Fig. 1d**). To gain insight into what pathways are perturbed in microglia by TIMP2 deletion, we performed over-representation analyses for gene ontology (GO)^31^ terms related to ‘biological processes’ and ‘cellular components’. Genes that were upregulated in KO microglia were significantly enriched in terms for “cell activation involved in immune response” and “positive regulation of cytokine production”, among others (**Fig. 1e**). Downregulated DEGs in KO microglia were significantly enriched in pathways related to “synapse” and “neurogenesis” (**Fig. 1e**). The transcriptomic profile of KO microglia may reflect a more activated and perhaps inflammatory state relative to WT microglia, potentially regulating their communication with neurons and ability to promote plasticity, as suggested in perturbed pathways identified by Ingenuity Pathway Analysis (IPA) (**Supplementary Fig. 1e**).

**Figure 1.**
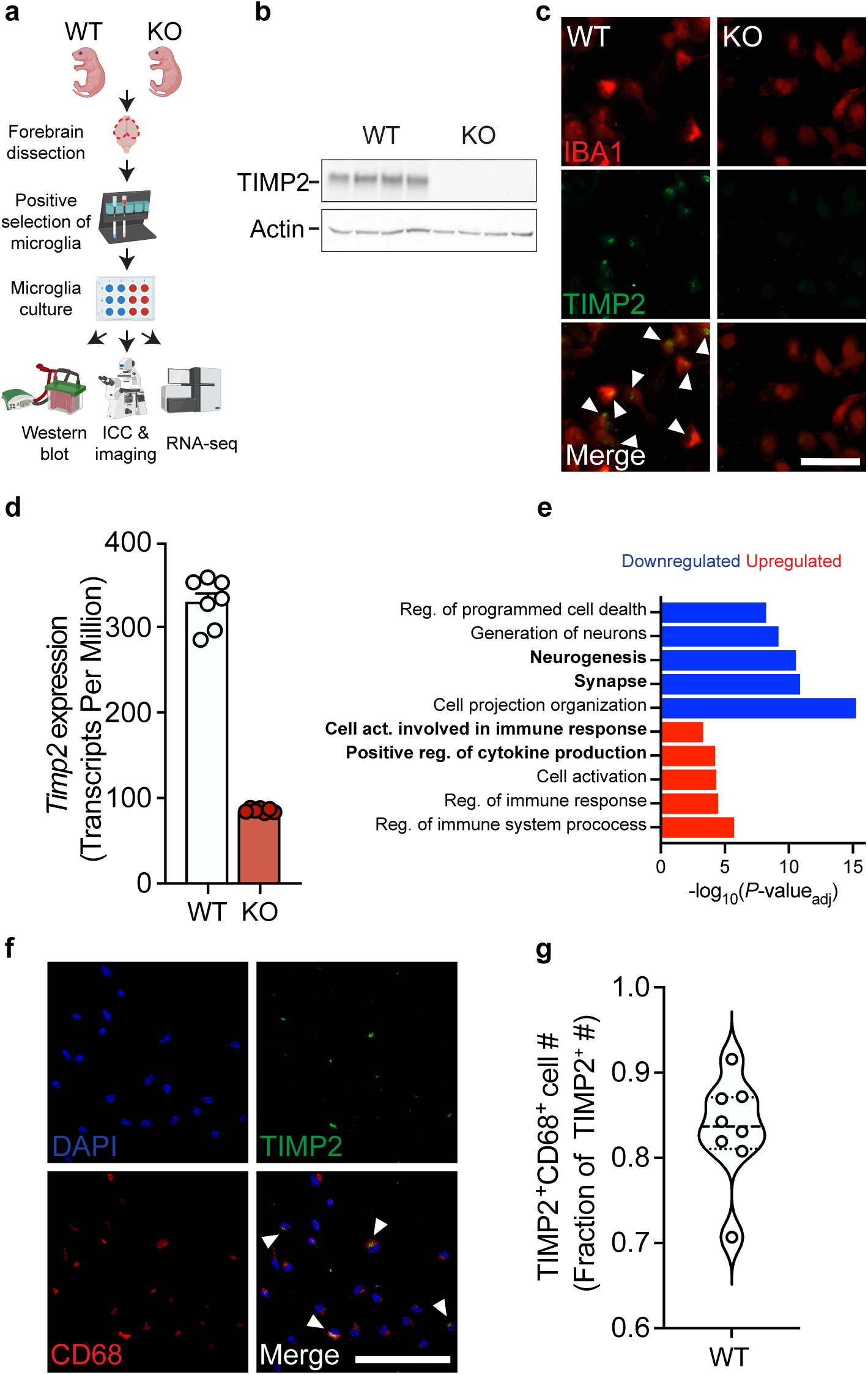
TIMP2 is expressed in microglia and its deletion leads to phenotypes associated with increased activation. **(a)** Schematic diagram of isolation, culturing, and assays performed on TIMP2 KO and WT primary microglia. **(b)** TIMP2 immunoblotting from primary microglia lysates isolated from TIMP2 WT and KO littermates (N = 4 samples per genotype, comprised of 5-6 pups (P5-P8) pooled per sample). **(c)** Representative images from cultures of TIMP2 KO and WT primary microglia (N = 3 pups (P6-P7) pooled per genotype; repeated in 2 additional experiments) stained with anti-TIMP2 and anti-IBA1 antibodies by ICC (scale bar = 100 μm). **(d)** Gene expression (transcript per million) values for *Timp2* from bulk RNAseq experiments from TIMP2 KO and WT primary microglia (N = 7 samples per genotype, with each sample representing a pool of 5 pups (P6-P7); sex-matched; mean ± SEM) with subsequent **(e)** over-representation analysis of GO terms (*P*_adj_ < 0.05) using upregulated and downregulated DEGs in TIMP2 KO vs. WT primary microglia. **(f)** Representative images from WT primary microglia cultures stained by ICC with DAPI, as well as anti-TIMP2 and anti-CD68 antibodies for co-localization analyses (N = 8 mice, 5-6-month-old mice; sex-matched; 2 independent experiments combined; scale bar = 100 μm), with **(g)** quantification of the fraction of TIMP2^+^ cells that are TIMP2^+^CD68^+^; violin plot of median with quartiles.

Given TIMP2’s potential relationship with activation state, we stained cultures of primary microglia isolated from adult mice and found that the majority of TIMP2-expressing cells co-express CD68, a lysosomal-associated protein found enriched in myeloid cells (**Fig. 1f-g**), pointing to a potential role for TIMP2 in regulating lysosomal function or activation state. While not all microglia were found to express TIMP2, a sizeable fraction of microglia expressing CD68 co-expressed TIMP2 (**Supplementary Fig. 1f-h**).

### TIMP2 deletion increases hippocampal microgliosis and extracellular inflammatory proteins

To evaluate the extent to which the putative pathways identified in our profiling of KO and WT microglia may reflect microglial phenotypes *in vivo* consistent with elevated activation state, we next sought to characterize microglial activation in KO and WT dentate gyrus of the adult hippocampus, a relevant region based on previous work demonstrating its role in regulating plasticity processes^16,25^. Using confocal microscopy, we evaluated microglia by co-staining for microglia marker IBA1 and activation marker CD68, which we previously found co-expressed with TIMP2 in primary microglia (**Fig. 1f-g**). As early as 3-4 months of age, KO mice exhibited a trend towards an increase in the percentage of dentate gyrus (DG) covered by IBA1 staining relative to WT (**Fig. 2a-b**), perhaps reflecting early changes in microglial morphology. While levels were modestly low given the early age, we detected significantly more microglia co-expressing CD68 in KO dentate gyrus compared to that from WT mice (**Fig. 2a,c**), as well as higher percentage area covered by CD68 (**Supplementary Fig. 2a-b**), indicating increased microgliosis. TIMP2 deletion did not affect the total number of microglia (**Supplementary Fig. 2a**,c).

**Figure 2.**
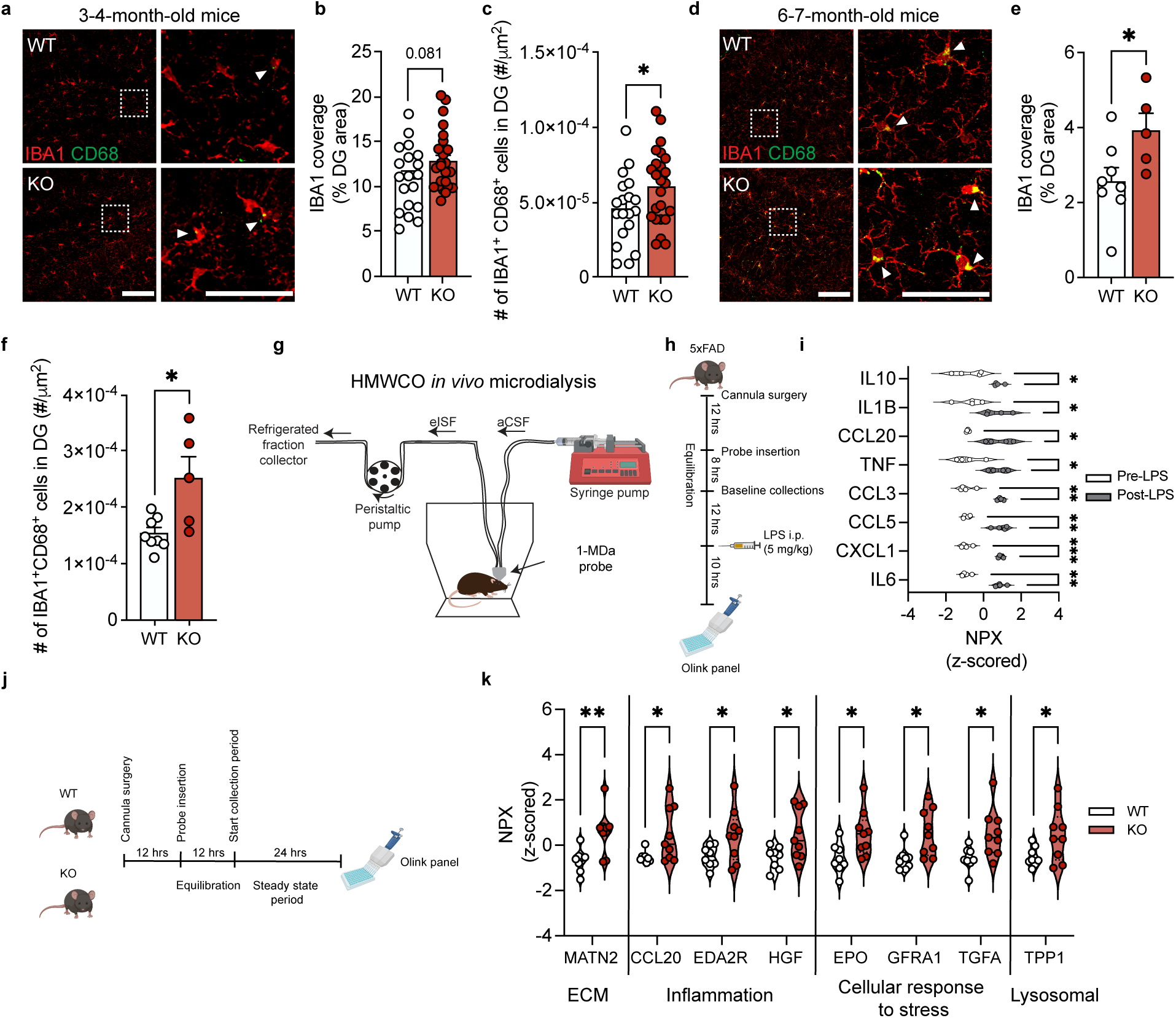
TIMP2 regulates microglial state and brain extracellular composition *in vivo*. **(a)** Representative confocal images of dentate gyrus (DG) of 3-4-month-old TIMP2 KO and WT mice stained with anti-IBA1 and anti-CD68 antibodies (left; scale bar = 100 μm) and magnified inset (right; scale bar = 50 μm) (N = 9-13 mice per group; sex-matched; arrowheads indicate IBA1^+^CD68^+^ cells) with corresponding **(b)** quantification of thresholded DG area covered by IBA1^+^ staining and **(c)** the number of IBA1^+^CD68^+^ cells normalized to DG area; mean ± SEM. **(d)** Representative confocal images of dentate gyrus (DG) of 6-7-month-old TIMP2 KO and WT mice stained with anti-IBA1 and anti-CD68 antibodies (left; scale bar = 100 μm) and magnified inset (right; scale bar = 50 μm) (N = 5-8 mice per group; female mice; arrowheads indicate IBA1^+^CD68^+^ cells) with corresponding **(e)** quantification of thresholded DG area covered by IBA1^+^ staining and **(f)** the number of IBA1^+^CD68^+^ cells normalized to DG area; mean ± SEM. **(g)** Schematic diagram of the high molecular-weight cut-off (HMWCO,1-MDa) *in vivo* microdialysis method to dialyze hippocampal ISF proteins and the corresponding **(h)** timeline of LPS treatment paradigm in mice undergoing microdialysis (N = 4; 6-7-month-old male 5XFAD mice) with **(i)** brain ISF protein measurements, represented as z-scored NPX values for significant proteins (paired t-test; \**P* < 0.05, \*\**P* < 0.01, \*\*\**P* < 0.001; median with quartiles). **(j)** Timeline of HMWCO *in vivo* microdialysis experiments to evaluate steady state levels of hippocampal ISF proteins in 2-3-month-old TIMP2 WT and KO mice (N = 8-9 mice per group; female mice) with corresponding **(k)** brain ISF protein measurements, represented as z-scored NPX values for significant proteins (unpaired t-test with Welch’s correction; \**P* < 0.05, \*\**P* < 0.01; median with quartiles). Student’s t test used in panels (b-c) and (e-f); \**P* < 0.05.

We next examined if these changes persist or worsen at a later age, given the well-established age-dependent microgliosis that occurs in the hippocampus^32^. At 6-7 months of age, we found that while the total number of microglia was not affected by TIMP2 deletion (**Supplementary Fig. 2d-e**), the percentage of DG area occupied by IBA1 immunoreactivity was significantly higher in the DG of TIMP2 KO compared to WT DG (**Fig. 2d-e**). To further characterize this change, we performed a morphology analysis that broadly categorizes microglia as “ramified”, “activated”, or “ameboid” based on their circularity using an established method (**Supplementary Fig. 2f**)^33–36^. Our morphology analysis revealed an increased number of morphologically “activated” microglia in the DG of KO mice relative to WT, with no change in the number of “ramified” or “ameboid” microglia (**Supplementary Fig. 2g-j**). At this age, we also detected a robust elevation in the number of IBA1^+^CD68^+^ cells in the DG of KO mice compared to WT, reflecting the increased state of activation (**Fig. 2d,f**).

In inflammatory contexts, it is well-established that microglia release cytokines and other factors into the extracellular space, affecting the cellular microenvironment of the brain. Given the increased microgliosis found in KO hippocampus, we next sought to assess whether deletion of TIMP2 alters the extracellular milieu. Conventional tissue-based assessment of cytokines and chemokines do not accurately reflect the composition of the brain interstitial fluid (ISF) given loss of compartmental barriers and effects of extraction methods on sensitivity, and this approach is limited to postmortem sampling, which may not sufficiently capture the dynamics of immune changes. To circumvent these issues, we employed high molecular-weight cut-off (HMWCO; 1-MDa) *in vivo* microdialysis to sample the hippocampal ISF of awake and freely moving mice (**Fig. 2g**)^37,38^. This technique has not yet been extensively applied to measure brain immune proteins, which are typically present in the brain ISF at very low levels. To establish this method’s potential to examine immune-related proteins in the brain ISF, we administered a single systemic injection of lipopolysaccardine (LPS; 5 mg/kg), an established pro-inflammatory stimulus^39^, as a proof-of-concept approach in a 5XFAD model of amyloid pathology that should exhibit high baseline cytokine and chemokine levels in the brain. Following microdialysis probe insertion into hippocampus and a subsequent equilibration period to allow levels of ISF proteins stabilize, as previously described^40,41^, samples were collected over a baseline period before LPS treatment. Samples were collected 10 hours following LPS and pre- and post-LPS period ISF samples were quantified using an exploratory proximity extension assay (PEA)-based panel of cytokines, chemokines, and related proteins (**Fig. 2h**). As shown in **Fig. 2i**, a number of expected pro-inflammatory cytokines and chemokines were significantly elevated in the hippocampal ISF following LPS treatment compared to the baseline period, including IL1β, CCL5, TNF, IL6, as well as anti-inflammatory protein IL-10, which has been shown to be released by microglia following LPS as a means to resolve the inflammatory response^42^ (**Fig. 2i**). Having successfully dialyzed and detected immune-related proteins using this method, we sought to assess the impact of TIMP2 deletion on the hippocampal extracellular milieu. 2-3-month-old WT and KO mice were subjected to the HMWCO *in vivo* microdialysis paradigm to sample hippocampal ISF during a stable, steady state period of 24 hours for measurement of proteins in pooled ISF samples (**Fig. 2j**). Interestingly, deletion of TIMP2 increased a number of proteins associated with inflammation in the hippocampal ISF (**Fig. 2k**), including chemokine CCL20 that we had found to be elevated following LPS stimulation (**Fig. 2i**), and EDA2R, a member of the tumor necrosis factor receptor superfamily whose levels correlates with increased age^43^. EDA2R cytokine signaling contributes to inflammation through activation of the NF-κB pathway. TIMP2 deletion also increased hippocampal ISF levels of HGF, a pleiotropic cytokine that modulates the inflammatory response through NF-κB signaling that is also elevated in CSF of AD subjects^44,45^. Several proteins known to be upregulated under neurotoxic conditions as a means to limit damage were also found to be upregulated in TIMP2 KO ISF, including EPO, GFRA1, TGFA^46,47^, as was the lysosomal serine protease TPP1 that is associated with phagocytic function^48,49^. As expected, we also found that loss of TIMP2 increases hippocampal ISF levels of Matrilin-2 (MATN2), an extracellular matrix (ECM) protein^25^ that signals through TLR4 to induce proinflammatory genes in macrophages to promote axonal damage^50^. Together, these changes in the hippocampal extracellular environment may reflect a more pro-inflammatory, debris-rich environment relative to that in WT hippocampus.

### Microglial TIMP2 regulates activation state

Our findings suggest that loss of TIMP2 alters microglial activation and contributes to an inflammatory extracellular environment, yet it remains unclear whether these differences are mediated by the pool of TIMP2 produced by microglia, consistent with its expression pattern demonstrated in our previous experiments (**Fig. 1b-c**). To explore the contribution of microglial sources of TIMP2 to the microglial phenotypes associated with global TIMP2 deletion, we used a TIMP2^fl/fl^ model we generated^25^ and crossbred them with *Cx3cr1*^CreERT2/+^ mice^51^ to inducibly delete TIMP2 in *Cx3cr1*-expressing cells (e.g., microglia). Using a previously established adult microglia isolation paradigm^52^ (**Fig. 3a**) that successfully yielded pure adult microglia in culture (**Supplementary Fig. 3a-c**), we first induced Cre-mediated recombination and then confirmed efficient deletion of microglial TIMP2 (**Supplementary Fig. 3d-e**). To examine how TIMP2 deletion in microglia affects canonical activation marker expression, we imaged primary adult microglia expressing CD68. Targeting microglial expression of TIMP2 significantly increased the number of cells expressing CD68 (**Fig. 3b-c**). We also quantified the number of primary microglia expressing senescent marker p16INK4a given the earlier pathway association in TIMP2 KO neonatal microglia (**Supplementary Fig. 1h**), which revealed a greater number of p16INK4a^+^ microglia from *Cx3cr1*^CreERT2/+^;*Timp2*^fl/fl^ mice compared to that observed from *Timp2^fl^*^/fl^ control mice. (**Fig. 3d-e**). We next examined the number of IBA1^+^CD68^+^ cells in the DG of 6-7 month-old *Cx3cr1*^CreERT2/+^;*Timp2*^fl/fl^ mice or *Timp2^fl^*^/fl^ control mice that had received tamoxifen (**Fig. 3f**). We observed an increase in the extent of IBA1 staining covering the DG and an increase in the number of activated microglia (IBA1^+^CD68^+^) (**Fig. 3f-h**). These findings suggest that removal of the microglial pool of TIMP2 leads to microgliosis phenotypes that are consistent with those observed in the setting of global TIMP2 deletion.

**Figure 3.**
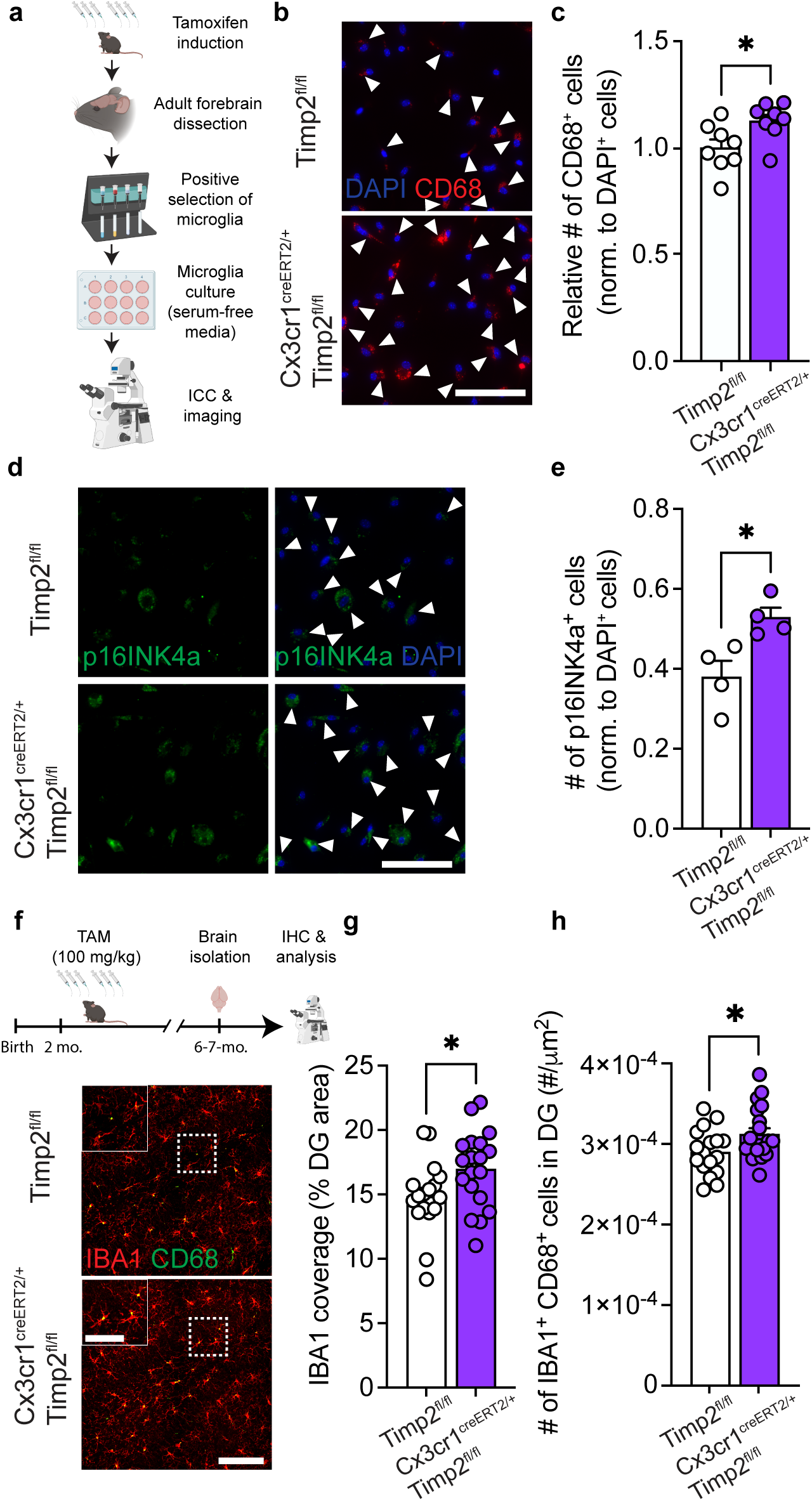
Removal of microglial TIMP2 increases states of activation and senescence. **(a)** Schematic diagram of tamoxifen-inducible microglial deletion of TIMP2 in mice and subsequent isolation of adult microglia, culturing, and imaging. **(b)** Representative images from primary microglia cultures of Timp2^fl/fl^ and Cx3cr1^CreERT2/+^; Timp2^fl/fl^ littermates stained by ICC with DAPI and anti-CD68 antibody (arrowheads indicate CD68^+^ cells; scale bar = 100 μm) with corresponding **(c)** quantification of the number of CD68^+^ cells normalized to total cell number (DAPI^+^) (N = 8 mice per group; two independent experiments combined; 5-6-month-old mice; sex-matched; mean ± SEM). **(d)** Representative images from primary microglia cultures of Timp2^fl/fl^ and Cx3cr1^CreERT2/+^; Timp2^fl/fl^ littermates stained by ICC with DAPI and anti-p16INK4a antibody (arrowheads indicate p16INK4a^+^ cells; scale bar = 100 μm) with corresponding **(e)** quantification of the number of p16INK4a^+^ cells normalized to total number of cells (DAPI^+^) (N = 4 mice per group, 7-month-old mice, sex-matched; mean ± SEM). **(f)** Timeline (top) of Cre induction paradigm with tamoxifen in Timp2^fl/fl^ control and Cx3cr1^CreERT2/+^;Timp2^fl/fl^ mice and downstream processing of tissue, with representative confocal images (bottom) of DG of 6-7-month-old Timp2^fl/fl^ and Cx3cr1^CreERT2/+^;Timp2^fl/fl^ mice stained with anti-IBA1 and anti-CD68 antibodies (scale bar = 100 μm) and magnified inset (scale bar = 50 μm) (N = 17-19 mice per group, sex-matched) with corresponding **(g)** quantification of DG area covered by anti-IBA1 staining and **(h)** the number of IBA1^+^CD68^+^ cells normalized to area. Mean ± SEM. \**P* < 0.05; Student’s t test.

### Microglia contribute to external pools of TIMP2 and its removal modifies myelin phagocytosis

TIMP2 is present at high levels in the extracellular space of the hippocampus^25^, likely reflecting contributions from various sources, including from neurons and microglia. Our data show that microglia express TIMP2 at both transcript and protein levels, yet it remains unclear whether microglia release TIMP2, where it can potentially alter microglial response to pathological debris, and whether targeting microglial TIMP2 affects associated phenotypes. To begin to explore these questions, we isolated and cultured adult WT microglia to examine TIMP2 release into the media across various timepoints in serum-free media, which was necessary given the high levels of endogenous TIMP2 present in plasma/serum, especially at early life stages^16^ (**Fig. 4a**). Immunoblotting analysis on media from these cultures revealed that microglia release TIMP2 in a time-dependent fashion (**Fig. 4b-c**). We then asked whether levels of TIMP2 released into the media is modulated by exposure to various biological substrates (zymosan, 10 μg/mL; LPS, 100 ng/mL), including those commonly encountered within the brain (myelin, 10 μg/mL). 12 hours following stimulation with these substrates, we measured TIMP2 levels in the conditioned media by immunoblotting. While zymosan stimulation did not change TIMP2 levels in the media relative to unstimulated conditions, LPS slightly increased levels, though this did not reach statistical significance, and myelin stimulation robustly increased (∼2-fold) TIMP2 release into the media relative to unstimulated conditions (**Fig. 4d-e**).

**Fig. 4.**
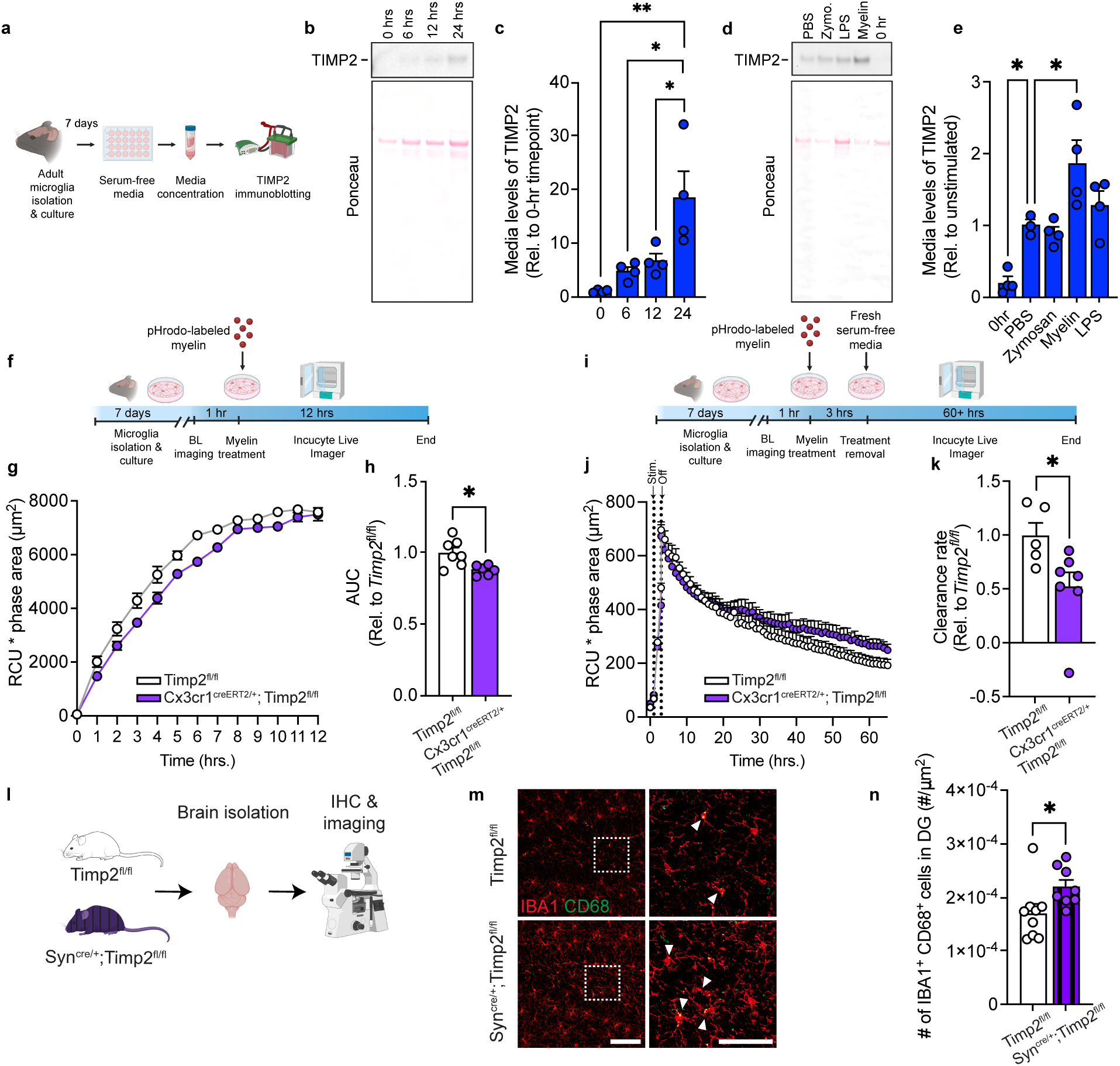
TIMP2 is released by microglia and alters microglial response to myelin. **(a)** Schematic diagram showing timeline of primary microglia culturing from adult mice and media assay for TIMP2 immunoblotting experiments. **(b)** TIMP2 immunoblotting (representative) from conditioned serum-free media from primary microglia isolated from 2-month-old WT mice, including Ponceau stain (bottom), with corresponding **(c)** TIMP2 band intensity quantification (N = 4 mice per timepoint; 4 independent experiments combined, with a full timecourse for each experiment; sex-matched; mean ± SEM). **(d)** TIMP2 immunoblotting (representative) from conditioned serum-free media from primary microglia isolated from 2-month-old WT mice at baseline (0 hours with no treatment) or 12 hours following treatment with either PBS, zymosan (10 mg/mL), myelin (10 mg/mL), or LPS (100 ng/mL), with corresponding **(e)** TIMP2 band intensity quantification (N = 3-4 mice per treatment; 4 independent experiments combined, with all treatments representing an experiment; sex-matched; mean ± SEM). **(f)** Schematic diagram of treatment of adult primary microglia with pHrodo-labeled myelin and subsequent Incucyte imaging to assess uptake. (**g**) Timecourse from an experiment with total integrated density (red object mean intensity [RCU, red, *x* average phase object area (cell, μm^2^)) of pHrodo-labeled myelin signal in primary microglia cultured from tamoxifen-treated 5-7-month-old Cx3cr1^CreERT2/+^;Timp2^fl/fl^ and Timp2^fl/fl^ control mice, with corresponding **(h)** uptake quantification using area under the curve (AUC; N = 6-7 mice per group; 2 independent experiments combined; sex-matched; mean ± SEM). **(i)** Schematic of experimental design to assess clearance of pHrodo-labeled myelin in Cx3cr1^CreERT2/+^;Timp2^fl/fl^ and Timp2^fl/fl^ control mice. (**j**) Timecourse from an experiment with total integrated density (RCU [red calibrated units]) *x* average phase object area (cell, μm^2^)) of pHrodo-labeled myelin signal in primary microglia cultured from 5-7-month-old tamoxifen-treated Cx3cr1^CreERT2/+^;Timp2^fl/fl^ and Timp2^fl/fl^ control mice, with corresponding **(k)** clearance rate quantification (from slopes of regressions), represented relative to Timp2^fl/fl^ control (N = 5-7 mice per group; 2 independent experiments combined; sex-matched; mean ± SEM). **(l)** Schematic diagram of neuron-specific TIMP2 deletion using *Timp2*^fl/fl^ and Syn^Cre/+^;*Timp2*^fl/fl^ mice for confocal imaging experiments. **(m)** Representative confocal images of DG from 2-3-month-old Timp2^fl/fl^ and Syn^Cre/+^;Timp2^fl/fl^ littermates stained with anti-IBA1 and anti-CD68 antibodies (left; scale bar = 100 μm) and magnified inset (right; scale bar = 50 μm; white arrowheads indicate IBA1^+^CD68^+^ cells) with corresponding **(n)** quantification of IBA1^+^CD68^+^ cell number normalized to DG area (N = 8-9 female mice per group; mean ± SEM). \**P* < 0.05, \*\**P* < 0.01; One-way ANOVA with Tukey’s post hoc test (c); One-way ANOVA with Dunnett’s post hoc test (e); Student’s t test (h,k,n).

Given that the brain accumulates abundant myelin debris with age^53^, the increase in TIMP2 in response to myelin stimulation motivated us to examine whether microglial TIMP2 modulates cellular response to myelin. We isolated adult microglia from tamoxifen-treated *Cx3cr1*^CreERT2/+^;*Timp2*^fl/fl^ mice and *Timp2*^fl/fl^ control mice and performed a myelin phagocytosis assay in which myelin was conjugated with pHrodo red, a pH-sensitive dye that fluoresces in the acidic endosomal/lysosomal compartment that was monitored over time using a live-cell imaging system (**Fig. 4f**)^54^. ‘Area under the curve’ analysis revealed significantly lower uptake of myelin in *Cx3cr1*^CreERT2/+^;*Timp2*^fl/fl^ microglia over a 12-hour period compared to the uptake observed in control microglia (**Fig. 4g-h**). We next assessed the rate of myelin clearance in the two groups by treating new cultures with myelin, followed by withdrawal of the treatment with fresh media to measure the rate of myelin elimination (**Fig. 4i**). We observed a lower rate of myelin clearance in *Cx3cr1*^CreERT2/+^;*Timp2*^fl/fl^ microglia relative to Timp2^fl/fl^ control microglia (**Fig. 4j-k**). Together, these results suggest that deletion of microglial TIMP2 impairs microglial phagocytosis of myelin.

We previously reported that neuron-derived TIMP2 released into the brain extracellular space is critical for hippocampal plasticity^25^. We next asked whether this additional extrinsic source from neurons can regulate microglial state by creating Syn^Cre/+^;*Timp2*^fl/fl^ and *Timp2*^fl/fl^ mice to target neuronal TIMP2 expression^25^ and then performing confocal imaging to evaluate microglial state using IBA1 and CD68 markers (**Fig. 4l**). Similar to our findings demonstrating that targeting the microglial pool of TIMP2 affects microglial state (**Fig. 3h**), we find that neuronal TIMP2 deletion significantly increases the number of IBA1^+^CD68^+^ cells in the DG (**Fig. 4m-n**), while the total number of microglia in this region was not affected by TIMP2 deletion (**Supplementary Fig. 4a-b**). Together, our findings indicate that microglia are responsive to modulation of extracellular levels of TIMP2, regardless of whether the source is from microglia or neurons.

### Systemic TIMP2 treatment reduces age-associated microgliosis and modulates microglial state

Given our data suggesting that extrinsic sources of TIMP2 regulate microglial state *in vivo*, we next asked whether extrinsic application of TIMP2 to aged primary microglia could directly affect microglial state phenotypes observed in our KO models, independent of regulation from other cell types. We first isolated microglia from 20-month-old WT mice and evaluated whether application of TIMP2 altered the number of IBA1^+^CD68^+^ cells in the culture (**Fig. 5a**). Treating aged primary microglia with TIMP2 reduced the number of CD68^+^ cells relative to control conditions (**Fig. 5b-c**), suggesting that TIMP2 can directly alter microglial state independent of its known effects on other CNS cell types^23,25,55^. Based on our previous results demonstrating that TIMP2 enters the brain from blood following systemic injection^16^, we next asked whether systemic treatments with TIMP2 in aged mice can regulate microglial state and function. Using the same dosing paradigm we previously showed improves learning and memory in aged mice (**Fig. 5d**)^16^, we injected aged WT mice systemically with TIMP2 (50 μg/kg), and isolated brains to evaluate changes in microglia after ∼2.5 weeks of injections. Systemic TIMP2 treatment reversed the consistent increase in microglial activation observed in our KO models, as reflected by the decreased number of IBA1^+^CD68^+^ cells in the DG of TIMP2-treated aged mice compared to vehicle-treated aged mice (**Fig. 5e-f**). We also detected a lower percentage of DG area occupied by CD68 staining, whereas the treatment did not affect the total number of IBA1^+^ cells (**Supplementary Fig. 5a-c**).

**Fig. 5.**
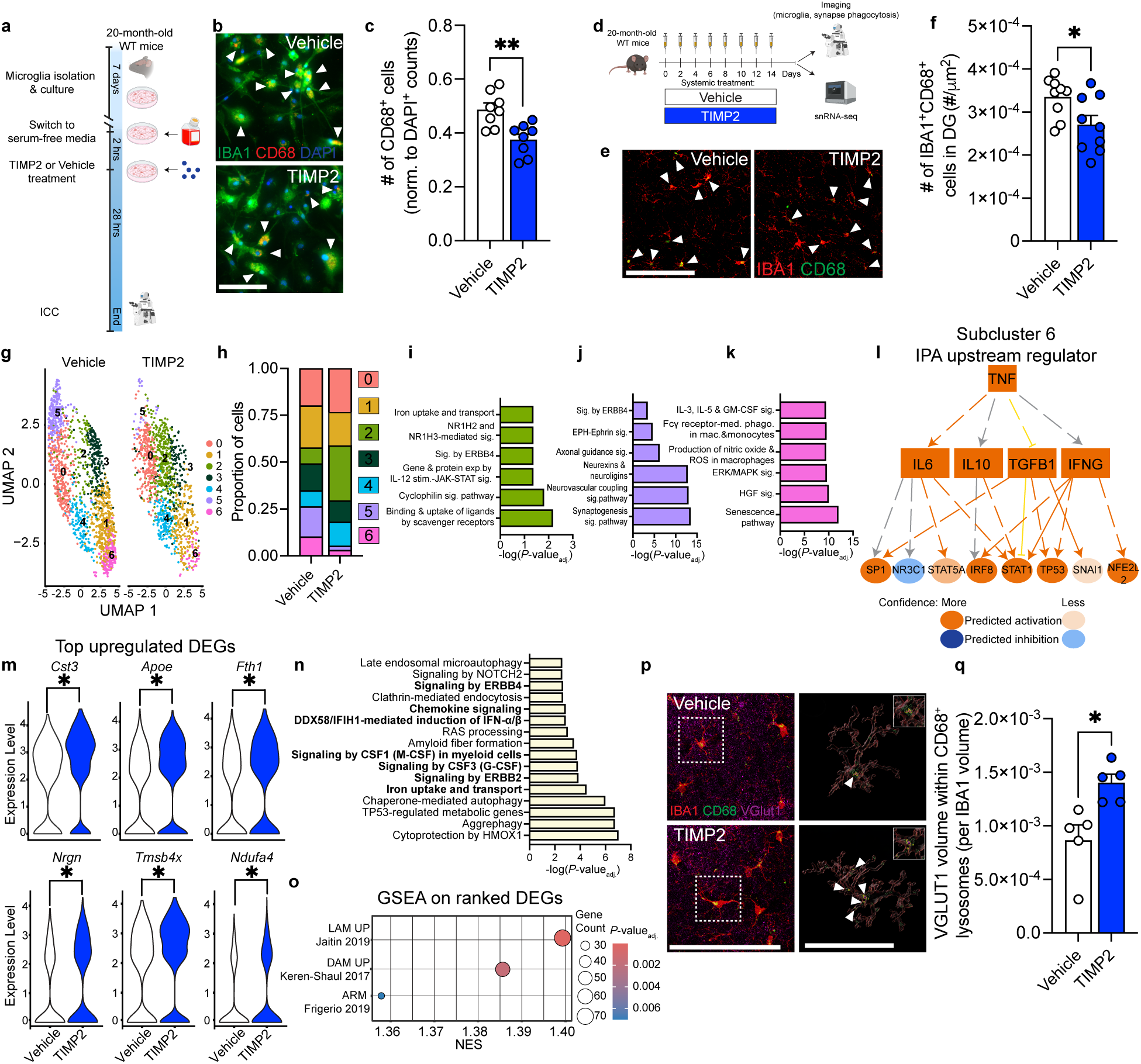
Youth-associated TIMP2 alters microglia in aged mice. **(a)** Schematic of experimental timeline of microglial isolation from 20-month-old WT mice, followed by incubation with TIMP2 or vehicle. (**b)** Representative images from cultures of WT primary microglia stained with anti-IBA1 and anti-CD68 antibodies by ICC (arrowheads indicate IBA1^+^CD68^+^ cells; scale bar = 100 μm), with corresponding **(c)** quantification of the number of CD68^+^ cells normalized to total cell number (N = 8 mice per treatment group, 2 independent experiments combined; sex-matched; mean ± SEM). **(d)** Schematic diagram showing paradigm of systemic treatment with recombinant TIMP2 or vehicle in 20-month-old WT mice, followed by downstream analyses. **(e)** Representative confocal images of DG from 20-month-old WT mice treated systemically with TIMP2 or vehicle, stained with anti-IBA1 and anti-CD68 antibodies (scale bar = 100 μm; white arrowheads indicate IBA1^+^CD68^+^ cells), with corresponding **(f)** quantification of number of IBA1^+^CD68^+^ cells normalized to DG area (N = 9 male mice per group; mean ± SEM). **(g)** Uniform Manifold Approximation and Projection (UMAP) plot of hippocampal microglial nuclei (2,831) isolated from 20-month-old WT mice treated systemically with vehicle or TIMP2 (N = 4 male mice per group combined as 2 samples per group). **(h)** Proportion of nuclei across microglial subclusters for each treatment group with corresponding **(i)** Significant canonical pathways from IPA for marker genes significantly upregulated in subcluster 2, **(j)** subcluster 5, and **(k)** subcluster 6. **(l)** IPA upstream regulator network based on subcluster 6 marker genes, with predicted relationships for target genes. **(m)** Violin plots of normalized gene expression values from selected top DEGs; \**P* < 0.0001, from DESeq2 analysis. **(n)** Significant canonical pathways from IPA on microglial DEGs (*P*-value_adj._< 0.05) from TIMP2-treated vs. vehicle-treated aged mice. **(o)** Dot plot from Gene Set Enrichment Analysis (GSEA) performed on ranked gene lists derived from differential expression analysis of microglia following TIMP2 treatment. Significantly enriched gene sets derived from published transcriptomic states are shown with corresponding Normalized Enrichment Score (NES), with point size representing number of overlapping genes and color indicating *P*-value_adj_. **(p)** Representative confocal images (left) of DG from 20-month-old WT mice treated systemically with TIMP2 or vehicle, stained with anti-IBA1, anti-CD68, and anti-Vglut1 antibodies (scale bar = 100 μm), with Imaris-based reconstruction (right) of insetted microglial volume (red), containing engulfed VGLUT1^+^ puncta (purple with yellow outlines indicated by white arrowheads) within microglial lysosomes (green) (scale bar = 50 μm). **(q)** Quantification of the volume of engulfed VGLUT1^+^ signal within CD68^+^ volume, normalized to microglia volume. Values represent N = 5 mice per group; mean ± SEM). \**P* < 0.05, \*\**P* < 0.01; Student’s t test.

To characterize how aged microglia respond to TIMP2 treatment at the transcriptomic level, we applied a single-nuclei RNA-sequencing (snRNA-seq) pipeline on hippocampi isolated from a new cohort of vehicle- and TIMP2-treated 20-month-old mice, based on an established method^56^. Following quality-control measures to remove ambient RNA, doublets, and poor-quality nuclei, we obtained 60,436 nuclei for downstream analysis in which nuclei from each sample were integrated and clustered using uniform manifold approximation and projection (UMAP) (**Supplementary Fig. 5d**). Combining use of ScType for automated cell-type identification^57^ and thorough interrogation of top marker genes for each cluster, we identified CNS cell types expected for the hippocampus (**Supplementary Fig. 5d-e**). We next subclustered cells identified as microglia (2,831 nuclei; **Fig. 5g**) and noted significant shifts in the proportion of cells in subclusters 2, 5, and 6 in mice treated systemically with TIMP2 compared to vehicle-treated mice (**Fig. 5h**; **Supplementary Fig. 5f**). To identify what processes characterize these TIMP2-shifted subclusters, we performed IPA on the significant genes defining these subclusters and examined expression levels of top marker genes. TIMP2 treatment increased the proportion of microglia in subcluster 2, a cluster characterized by the highest expression of *Apoe,* a gene that regulates phagocytosis and lipid uptake^28,58^ (**Supplementary Fig. 5g**), as well as by iron handling (“uptake and transport”) and immune response processes (“cyclophilin signaling”) (**Fig. 5i**). TIMP2 treatment shifted subcluster 1, albeit modestly (**Supplementary Fig. 5f**), which was characterized by genes relating to phagosome formation and signaling in myeloid cells (**Supplementary Fig. 5h**). TIMP2 treatment significantly reduced the proportion of microglia in subcluster 5 (**Fig. 5h; Supplementary Fig. 5f**), a cluster defined by “axon guidance signaling”, “synaptogenesis signaling”, and “neurexins and neuroligins” (**Fig. 5j**). This subcluster, along with 2 other microglial subclusters with neuron-associated genes, also had highly enriched expression of *Gria1* (**Supplementary Fig. 5g**), a gene linked to microglial-neuron interactions^59,60^. Intriguingly, we found that TIMP2 treatment resulted in a significant decrease in proportion of subcluster 6 microglia (**Fig. 5h; Supplementary Fig. 5g**), which were defined by pathways associated with production of “nitric oxide and reactive oxygen species (ROS) in macrophages”, “senescence”, “Fcγ receptor-mediated phagocytosis”, “IL-3, IL-5, and GM-CSF signaling”, and “HGF signaling” (**Fig. 5k**). Further analysis of subcluster 6 markers revealed “TNF” as a top upstream regulator (**Fig. 5l**), activating proinflammatory^61–63^ and senescence-associated^64^ pathways through IL6, IFNG, STAT1/STAT5A, IRF8, and TP53, as well as IL10 as a counter-regulatory response protein^65^, together suggesting that TIMP2 treatment of aged mice reduces the proportion of microglia associated with a pro-inflammatory and senescent-like state.

Given these shifts in microglial subclusters following TIMP2 treatment, we next aimed to compare overall microglial gene expression changes following TIMP2 treatment. We found that TIMP2 treatment resulted in differentially expressed genes (DEGs; 196 upregulated, 68 downregulated) in the overall microglia cluster. *Cst3* and *Apoe* were the two most significantly upregulated genes (**Fig. 5m**). *Apoe* is a well-characterized regulator of the DAM transcriptional state^28^ and is upregulated in microglia phagocytosing apoptotic neurons^58^, while *Cst3* is localized in the lysosome, where it regulates lysosomal homeostasis^66^. We also observed a change in ferroptosis-associated *Fth1*^67^ and genes linked to metabolism and inflammation, including *Ndufa4*^68^ and *Tmsb4x,* which negatively regulates NFκB signaling^69^ (**Fig. 5m**). Pathway analysis on the overall DEGs reflected changes in cytokine/chemokine and interferon signaling, as well as altered uptake processes (ferroptosis), and potential changes in microglia-neuron interactions (ERBB4 and ERBB2 signaling) (**Fig. 5n**). We performed pre-ranked Geneset enrichment analysis (GSEA)^70^ to determine if genes, ranked by signed fold-change*-log_10_*P*-value_adj_, were enriched within gene sets previously associated with different microglial states, including disease-associated microglia (DAM)^28^, lipid-associated macrophages (LAM)^71^, which was identified as a protective state in adipose tissue of mice on a high-fat diet^71^, lipid-droplet-accumulating microglia (LDAM)^12^, activated response microglia (ARM)^72^, interferon-related^73^ and homeostatic microglia^72^. Interestingly, we found that TIMP2 treatment resulted in positive enrichment in LAM, DAM, and ARM gene sets in microglia (**Fig. 5o; Supplementary Fig. 5i**), supporting a putative enrichment in genes associated with protective response to damage.

### Systemic TIMP2 restores phagocytic capacity in aged microglia

Phagocytic capacity of microglia is diminished in aging and in proinflammatory states^8,9,12,74^, and targeting this process may facilitate effective clearance of dead cells or debris^9,10,75^. Given the TIMP2-induced changes observed in microglial phagocytosis-related pathways and in putative microglial-synapse interactions in snRNA-seq experiments, we evaluated how systemic TIMP2 treatment affects the ability of microglia to phagocytose synaptic material in the DG. We hypothesized that TIMP2 treatment elevates microglial phagocytosis in aged DG, especially given our results showing that microglia-specific *Timp2* deletion reduced myelin phagocytosis (**Fig. 4g-h**). To evaluate microglial phagocytosis in aged mice following TIMP2 or vehicle treatment (**Fig. 5d**), we performed confocal imaging of sections stained with antibodies for IBA1, CD68, and synaptic marker VGLUT1, followed by Imaris-3D reconstruction to visualize lysosome-associated synaptic puncta in individual microglia. We found that TIMP2 treatment increased the volume of VGLUT1^+^ puncta within microglia lysosomes in the DG relative to vehicle treatment (**Fig. 5p-q**), reflecting enhanced phagocytic function of a process typically diminished with aging^8,12^.

## Discussion

In the current study, we define a novel role of youth-associated protein TIMP2 in regulating microglial state in the context of aging pathology and in normal brain homeostasis. Global, microglial-specific, and neuronal-specific deletion of TIMP2 accelerated activation and associated deleterious phenotypes of microglia in the hippocampus. TIMP2 also appeared to modulate the phagocytic uptake and clearance of physiological substrates by microglia, including myelin, while regulating the inflammatory state of these cells. Given that various sources of expression contribute to the brain extracellular pool of TIMP2, we explored how systemic supplementation with TIMP2 in aged mice affects microglia, which reversed many of the phenotypes associated with TIMP2 removal, including microgliosis and impaired phagocytosis, while highlighting transcriptomic shifts in microglia associated with deleterious microglial phenotypes. Together, our results highlight that microglia are sensitive to extracellular levels of TIMP2, and that increasing TIMP2 promotes a shift toward a more protective state that may counteract age-associated pathology.

Early studies provided intriguing evidence that exposure to the systemic environment may affect microglia^18,76^ and that these cells can take up proteins originating in plasma^20,21^, yet little is known of the effects of such proteins on microglia. Specifically, recent work found that labeled plasma proteins are found within microglia of healthy brains in a process that appears to involve a specialized subset of microglia characterized by antigen presentation, as well as high metabolic and phagocytic activity^21^, though it remains unclear whether these processes diverge in the aged brain. A further recent study identified PF4 as a platelet factor that modulates microglial state, though it may not penetrate the brain appreciably and instead acts on peripheral T cells to indirectly modulate the CNS^18,77^, highlighting that microglia likely respond to blood-borne factors through a variety of mechanisms. In the case of TIMP2, previous work found that labeled TIMP2 enters the brain after peripheral delivery in aged mice^16^. Our current data examining the impact of neuronal, microglial, and systemic sources of TIMP2 on microglial phenotypes collectively support the concept that extracellular pools of TIMP2 regulate microglial function.

A growing literature has highlighted roles for TIMP2 in regulating overall hippocampal function by modulating synaptic plasticity mechanisms^16,25^, raising the possibility that some of the effects observed in microglia may reflect, in part, its action on neurons. While this remains a possibility, we find that aged primary microglia treated with TIMP2 recapitulate phenotypes observed in the intact brains of aged mice treated systemically with TIMP2, suggesting that microglia are directly responsive to TIMP2. In terms of its molecular targets, a recent study using a modified TIMP2 construct lacking MMP inhibitory activity was still sufficient to enhance hippocampal plasticity phenotypes, arguing for non-canonical targets for TIMP2 activity^23^ beyond classical interactions with MMPs. Given the many distinct interactions for TIMP2 activity that have been identified^24^, future studies should focus on delineating molecular targets for specific CNS functions.

To further examine how TIMP2 modulates microglial state, we applied a combination of *in vivo* and *in vitro* approaches across multiple ages, which revealed a consistent phenotype of heightened microglial activation in the setting of TIMP2 deficiency that is reversed with TIMP2 supplementation. To characterize the local extracellular environment, we implemented an *in vivo* microdialysis method that permitted sensitive assessment of immune-related proteins, an approach that revealed marked alterations in the extracellular milieu in factors related to inflammation, lysosomal activity, and cellular stress responses, reflecting altered neuroinflammatory environment. Our transcriptomic analysis of aged microglia from TIMP2-treated mice largely corroborated these roles of TIMP2 as a driver of protective microglial states. In microglia of aged mice treated with TIMP2, we find enrichments in sets of genes resembling the LAM^71^, DAM^28^, and ARM^72^ states associated with protective macrophage/microglia function. DAM and ARM states are both associated with protective responses to aging and AD pathology^28,72^ while being characterized by elevated *Apoe* expression, a top upregulated gene in TIMP2-treated microglia in our dataset. Several studies have demonstrated that these states, and induction of genes associated with these states, are necessary for microglia to appropriately respond to damage^28,58,72,78,79^. We also found significant enrichment in the LAM geneset in TIMP2-treated microglia, which has been argued to be a protective myeloid state in adipose tissue of mice on a high-fat diet^71^. These macrophages are involved in phagocytosis, lipid catabolism, and energy metabolism^71^, largely fitting with our observed upregulation of genes related to phagocytosis, metabolism, and modulation of inflammatory response in microglia from TIMP2-treated mice. The enrichment in genes associated with LAM, DAM, and ARM states with TIMP2 treatment may reflect a shared change in microglial response to lipid-rich damage, including the myelin debris encountered by aged microglia, possibly a protective response to aging that is bolstered with TIMP2 treatment. Corroborating the protective effects, we see a corresponding loss of microglial subpopulations characterized by deleterious pathways in microglia from TIMP2-treated mice, including senescence, production of nitric oxide and ROS, and pro-inflammatory programs driven by TNF. A previous *in vitro* study using the BV2 cell line suggested that TIMP2 may have anti-inflammatory roles, including production of IL10 and reduction of pro-inflammatory cytokines following LPS stimulation^80^, supporting our *in vivo* observations.

A consistent phenotype we observed across experiments modulating TIMP2 levels is the shift in microglial activation and associated lysosomal and phagocytic capacity. Using lysosomal-associated marker CD68, we show that removing TIMP2 increases activation state, while reducing uptake and removal of myelin, together supporting that TIMP2 regulates how physiological material is removed from the extracellular space. Interestingly, by sampling the brain extracellular space of mice lacking TIMP2, we detected elevations in the lysosomal serine protease TPP1, perhaps reflecting response to aberrant debris removal by microglia, a phenomenon observed under conditions of lysosomal stress^81,82^. Conversely, TIMP2 treatment of aged mice reduces microglial activation and increases phagocytic capacity—a function typically diminished with age and under proinflammatory conditions^8,9,74^. These data dovetail with our transcriptomic data from treated aged mice that reveal upregulation of genes involved in phagocytic processes and metabolism and a shift away from proinflammatory microglial states, perhaps reflecting a more efficient endolysosomal capacity. In addition to the benefit of clearing debris to protect the local brain environment, the process of phagocytosis itself can also inhibit proinflammatory responses^83^, though future work will need to identify the directionality of these changes following treatment with TIMP2 and related factors. Together, this work suggests that TIMP2 modulates the ability of microglia to remove extracellular material, enabling them to more effectively engage in phagocytosis while reducing the inflammatory environment in the setting of age-related pathology.

Our findings demonstrate that TIMP2 modulates microglial state, enhancing phagocytic function and dampening inflammatory activation in the aged brain. By promoting a more protective set of microglial phenotypes, TIMP2 may help counteract accumulation of debris and chronic inflammation that characterize the aged CNS environment. Given that genetic risk for AD is enriched in genes involved in microglial phagocytosis and endolysosomal function^84^, future studies should explore how TIMP2 influences these pathways in contexts of aging and age-related neurodegenerative conditions. Together, our work positions TIMP2 as a promising candidate for mitigating microglial dysfunction and preserving brain health across the lifespan.

## Materials and Methods

### Animals

Animal procedures, care, and handling were performed in accordance with the NIH Guide for Care and Use of Laboratory Animals and the Icahn School of Medicine at Mount Sinai Institutional Animal Care and Use Committee. All mice were maintained on C57Bl/6 background. TIMP2 −/− (KO) mice were purchased from Jackson Laboratory, and 5xFAD mice were purchased from MMRRC and subsequently maintained in-house alongside WT littermates. *Timp2*^fl/fl^ mice were generated as previously described^25^ and crossbred with *Cx3cr1*^CreERT2/+*Lit51t*^ mice or Syn^Cre/+^ lines (Jackson) to permit microglial and neuronal deletion^25^ of TIMP2, respectively. To induce conditional deletion of microglial *Timp2*, 2-month-old mice were administered 100 mg/kg tamoxifen (Sigma Aldrich Fine Chemicals Biosciences, T5648) in corn oil (Sigma, C8267) 5 times by oral gavage, with at least 48-h separation between gavages^78^. Cre-negative *Timp2*^fl/fl^ littermates were also given tamoxifen as controls. Aged WT C57BL/6 mice were obtained from the National Institute on Aging aged rodent colony. Male and female mice were used unless otherwise indicated.

### Neonatal microglia isolation

Using an established protocol^85^, we dissected forebrain from pups (P5-P8) and manually homogenized the tissue using a dounce homogenizer (VWR, KT885300-0002), prior to debris removal using a 20% Percoll gradient (GE Healthcare, 17-0891-02). Microglia were positively selected using CD11b microbeads (Miltenyi Biotec, 130-049-601), LS columns (Miltenyi Biotec, 130-042-401), and a QuadroMACS Separator according to manufacturer instructions. For western blot analysis, cell pellets were collected and frozen at −80C until use. For ICC and bulk RNAseq experiments, cells were manually counted with a hemacytometer and Trypan blue staining to assess cell viability. Cells were plated at a density of 200,000 microglia/well in 96-well poly-d-lysine-coated plates (Corning, 354461) in Dulbecco’s modified Eagle’s medium (DMEM) supplemented with 10% FBS (Sigma, F4135, Sigma) and 1% penicillin–streptomycin (Gibco, 15140). Microglia recovered for a minimum of 2 days in tissue culture incubator before experiments.

### Adult microglia isolation

Following an established protocol^52^, mice were perfused with ice-cold PBS, cerebellum and olfactory bulbs were removed, and remaining brain tissue was enzymatically dissociated using the Neural Tissue Dissociation Kit (P) (Miltenyi Biotech, 130-092-628) according to manufacturer instructions. During incubation in enzymatic buffer, large, medium, and small fire polished Pasteur pipettes were used to mechanically dissociate tissue. Following filtration and centrifugation, samples were incubated with CD11b microbeads (Miltenyi Biotec 130-049-601) and subsequently separated using LS columns (Miltenyi Biotec 130-042-401) and QuadroMACS Separator using manufacturer’s instructions. Cells were plated on imaging plates (Cellvis, P96-1.5H-N) or poly-L-Lysine-coated glass coverslips (Electron Microscopy Sciences, 72292-04) in DMEM supplemented with 10% FBS and 1% penicillin–streptomycin. Microglia were allowed to recover for 7 days in a tissue culture incubator before experiments. Wells were excluded from analysis if cultures did not meet viability or quality control criteria (e.g. excessive debris).

### *In vitro* phagocytosis assay

Following a 7-day recovery period post-isolation, conditioned media from primary microglial cultures was removed and replaced with serum-free media before measurements in the IncuCyte S3 live-cell analysis system (Sartorius) that contained a tissue culture incubator at 37C/5% CO2 for baseline imaging. Cells were treated with pHrodo-labeled myelin^54^ (10 μg/mL), generously provided by the laboratory of Dr. Alison Goate, and imaged hourly at 20x using phase contrast and red fluorescent signal. For myelin clearance experiments, following a 3-hour incubation with pHrodo-labeled myelin, cells were washed with PBS and fresh serum-free media was added that did not contain myelin. Total integrated density was calculated by multiplying the average red object mean intensity (RCU, red calibrated units) by the average phase object area (cell, μm^2^) and plotted over time^86^. For both uptake and clearance experiments, two independent experiments were conducted with 3 technical replicates per animal that were averaged.

### *In vitro* microglial stimulation

To assess TIMP2 secretion with various substrates, cultures of adult WT microglia were stimulated with various substrates in serum-free media at the following concentrations: Zymosan 10 μg/mL, myelin 10 μg/mL, LPS 100 ng/mL, or PBS control. 12 hours following stimulation, conditioned media was collected and concentrated for immunoblotting. For experiments examining activation of primary microglia from aged mice following TIMP2 treatment, cell media was replaced with serum-free media 2 hours prior to incubation with TIMP2 (2 μg/mL, R&D Systems, 6304-TM-010) or vehicle (PBS) for 28 hours, after which cells were fixed and stained to evaluate markers.

### Bulk RNA-sequencing of neonatal microglia

Forebrains from neonatal mice were pooled by genotype, and then microglia were isolated and plated in 12-well poly-d-lysine-coated plates (Corning, 354470) at a density of 600,000 cells per well, 2 wells per genotype. After 72 hours, trypsin was used to remove cells prior to centrifugation for 5 minutes at 1230 x g, followed by resuspension in PBS, and re-centrifugation. Pellets were stored at −80C before further processing and sequencing by Genewiz (Azenta) with each sample representing independent isolations from pooled pups. RNA was extracted and quality was measured using TapeStation Analysis Software 3.2 (Agilent), and concentration was determined using Qubit assay, with all samples exhibiting RNA Integrity Number (RIN) = 10. Sequencing libraries were prepared with poly(A) selection and sequenced using Illumina Hiseq (2×150bp paired-end) according to manufacturer protocols. Reads were mapped to the Mus musculus GRCm38 genome. Differential expression was performed using DESeq2, and differentially expressed genes (*P*<0.05) were used to perform Ingenuity Pathway Analysis (IPA, Qiagen), as well as gene ontology over-representation analysis with the MSigDB^31^ database on significantly upregulated and downregulated genes separately.

### Immunoblotting

To perform western blot analysis of TIMP2 in primary microglia, cell pellets from pooled samples were lysed in RIPA buffer (Thermo Fisher) containing protease Inhibitor cocktail (Roche) for 30 min on ice, collecting lysates following a 30-min centrifugation at 15,000 x g. 80 μg of protein per sample was loaded on Bolt 4-12% Bis-Tris Plus Gels (Invitrogen). TIMP2 from conditioned media was spun at 1,000 x g for 10 min to remove debris. Protease inhibitor cocktail was added to supernatant prior to concentration of media by centrifugation at 4,000 x g for 40 min using Amicon Ultra-15 centrifugal filter unit (Amicon, UFC901024) before loading on 4-12% NuPAGE Bis-Tris denaturing gels (Invitrogen). Gels were transferred, the blots probed with anti-TIMP2 (1:5000; D18B7, Cell Signaling) or anti-actin (1:10000; A5060; Sigma), and then developed as previously described^25^, with multiple blots per experiment developed in parallel where necessary. Band intensities were quantified using ImageJ software as described^41^.

### Immunocytochemistry

Following fixation with 4% paraformaldehyde for 20 min at room temperature, primary microglia were washed three times with PBS, incubated with blocking solution (10% donkey serum and 0.1% triton-X in PBS) for 1 hour, and then incubated overnight at 4C with primary antibodies in blocking solution at the following concentrations: IBA1, 1:500, (019-19741, Wako); CD68, 1:200, (MCA1957, BioRad), TIMP2, 1:100 (AF971, R&D systems, [2-day incubation]), p16INK4a, 1:500, (MA5-17142, Invitrogen). Cells were then washed and incubated in their corresponding secondary antibody for 1 hour. Staining was visualized and imaged using the Keyence BZ-X700 microscope or Zeiss LSM780 upright confocal microscope using a 40x objective. 2-3 wells per mouse were plated and 3-4 images per well were averaged for 3 technical replicates per mouse. The value for each mouse represents the mean of these replicates.

### Immunohistochemistry and imaging

Following anesthetization with a ketamine (90 mg/kg) and xylazine (10 mg/kg) cocktail, mice were perfused transcardially with ice-cold 0.9% saline. Brains were dissected and postfixed in 4% paraformaldehyde for 48 hours and then preserved with 30% sucrose in PBS. 40-μm sections were prepared from hemibrains and sectioned using a freezing-sliding microtome and stored at −20C in cryoprotectant media prior to immunohistochemistry according to our previous protocol^16^. Briefly, free-floating sections were blocked in appropriate serum (10%) before incubation in primary antibody at 4C overnight. Primary antibodies were used at the following concentrations: IBA1, 1:500, (Wako, 019-19741); CD68, 1:200, (MCA1957, BioRad); VGLUT1, 1:2000, (135318, Synaptic Systems); CD68, 1:500, (137001, BioLegend, paired with IBA1 and VGLUT1 co-staining). Sections were then incubated in the corresponding fluorescently-conjugated secondary antibodies for 1 hour (1:200 in TBST), followed by DAPI, where indicated, for 15 min. Stained sections were mounted and coverslipped with ProLong Gold (Invitrogen).

To image staining in the dentate gyrus, 2×2 tile-scanning images were acquired through the entire z-plane of sections (1-μm intervals across ∼40μm) at 40x/1.4 oil DIC objective using a Zeiss LSM780 upright confocal microscope. For all imaging experiments, 3-4 sections per animal were imaged according to stereological principles, as previously described^16^. FIJI was used to perform thresholded “percentage area” or count analysis of cells expressing CD68 and/or IBA1 across mouse experiments in a blinded fashion. Count data was normalized to the area of the analyzed ROI.

#### Imaris-based imaging and analysis

Following IHC for VGLUT1, IBA1, and CD68, images containing dentate gyrus were acquired using a 63X oil-immersion objective lens on a Zeiss LSM780 with 0.33-μm z-step size across ∼20-μm z-stack (1024×1024 pixels). 3D reconstruction of microglia was performed using Imaris (Imaris 10.2, Oxford Instruments). Microglial surfaces were created using machine learning-based training on the IBA1 channel, which were then used to mask the CD68 channel. A surface was then created for the microglial-masked CD68 and used to mask VGLUT1 signal. A surface for VGLUT1 signal within a microglial lysosome was created, together creating VGLUT1^+^ surfaces within a microglial lysosome. The total volume of VGLUT1 within lysosomes was normalized to the volume of the microglial surface analyzed. Three sections per mouse were imaged, and results for microglia were averaged to represent a value for each mouse. A total of 44-47 microglia were analyzed per group.

#### Microglia morphology analysis

We employed an established protocol defining microglia morphologies based on their circularity ((4π Area) / Perimeter^2^) assessed by FIJI, where values approaching a value of 1 approximate a circular morphology^33–36^. “Ramified” morphology was categorized according to circularity between 0 and 0.400, “activated/reactive” microglia are defined by circularity of 0.500-0.699, and “ameboid” morphology are defined by circularity of 0.700-1.000. Images were imported to FIJI, converted to 8-bit, background-corrected (rolling ball radius of 50 pixels and despeckled) equally across images/groups, and then classified into the three morphology types according to circularity. The number of microglia corresponding to each morphology class was normalized to stereologically-defined dentate gyrus areas.

### Recombinant protein injections

For experiments in which mice were treated systemically with TIMP2, aged mice were given intraperitoneal (i.p.) injections every other day with either vehicle (PBS) or recombinant mouse TIMP2 (R&D Systems, 6304-TM-010) for a total of 8 injections, using our established protocol, a method used previously to provide revitalization of hippocampal plasticity and cognitive function^16^.

### Nuclei Isolation and snRNA-seq

Full hippocampi were isolated, flash-frozen, and stored at −80C until processing. Hippocampi were thawed on ice, and nuclei were isolated using established protocols^56^. Briefly, each hippocampi was mechanically dissociated using a dounce homogenizer (VWR) with 2 ml of EZ lysis buffer (Millipore Sigma). Samples were centrifuged, resuspended in fresh lysis buffer, and centrifuged again. Cell pellets were gently washed with 4 ml ice-cold PBS, and supernatant were removed prior to resuspending pellets in 100 μl of ice-cold PBS with 0.04% BSA (NEB) and 1 U/μL RNAse inhibitor (Millipore Sigma). Each sample was filtered through 40-μm FlowMi cell strainers (Millipore), and nuclei were counted (aided by trypan blue) to assess nuclei viability using an EVOS M7000. Two biological samples from the same treatment condition were pooled to load 20,000 nuclei total (10,000 nuclei each) for a total N=2 samples per treatment condition. Libraries were prepared and sequenced by the Single-cell & Spatial Technologies Team, Center for Advanced Genomics Technology Genomics Core (ISMMS) using the 10x Genomics Chromium platform for library preparation (V2 3’ GEX protocol) according to manufacturer’s instructions. Briefly, cDNA was extracted from Gel-Bead in Emulsions (GEMs) obtained from the sample chip in the Chromium controller, and library quality control was performed with MiSeq Nano. Libraries were run on Illumina NovaSeq 6000 S4 Flowcell (2×100 nt paired-end read length) configuration, targeting 50,000 reads per cell.

Raw read data was analyzed by 10x Cell Ranger (v7.1.0) to align to reference genome (Mus musculus, version mm10 2020-A). Downstream processing was performed in the R programming environment (v4.2.0 or v4.4.2). Ambient RNA was removed from samples using SoupX^87^ and manual application of a stringent contamination fraction on a per sample basis. After initial filtering, doublets were identified and removed using DoubletFinder^88^. Further filtering involved removing genes expressed in fewer than 3 cells, cells with a UMI count under 500, and cells with only 500 genes per cell to ensure we analyzed high-quality cells. The Seurat object was then log-normalized, and variable features were identified using the FindVariableFeatures function. ScaleData was then used to regress cells with more than 2% mitochondrial, ribosomal large subunit (RPL) and ribosomal small subunit (RPS) gene expression content. Using STACAS, a semi-supervised data integration method, we integrated all samples^89^, which was used for cell clustering. Using Seurat (v5.2.1), integrated samples were scaled, and clusters were identified using FindNeighbors (using top 50 principal components) and FindClusters (Resolution=0.25). To identify markers of each cluster, the RNA assay was processed using NormalizeData, FindVariableFeatures, ScaleData (regressing out > 2% mitochondria, RPS, and RPL), and FindMarkers. We determined cluster identity by assessing the top 30 markers on DropViz, further confirming identities with ScType^57^ and by examining literature-based expression of marker genes for each cell type. Clusters identified to be microglia were subclustered (using the top 15 principal components and a resolution of 0.5). Clusters suspected as containing doublets, i.e., marked by high nCount_RNA and nFeature_RNA, as well as a lack of microglial genes as cluster markers, were removed, and the remaining cells were re-clustered using the top 15 principal components and a resolution of 1. Pathways representing each subcluster were evaluated using the significantly upregulated marker genes in each subcluster (*P*_adj_ < 0.05; via FindAllMarkers) and by running these markers in Ingenuity Pathway Analysis (IPA, Qiagen) and gene ontology over-representation analysis using MSigDB^31^. To compare shifts in the proportion of subclustered cells between treatment conditions, we employed scProportionTest^90^.

Differential gene expression analysis for the microglia cluster was conducted using DESeq2, and IPA was run on significant genes (*P*_adj_ < 0.05). After removing genes not mapping to Entrez identifiers, we performed GSEA using a ranked list of all genes tested in the differential expression analysis to assess enrichment in published gene sets representing characterized microglial states^70^. GSEA was performed in R using the clusterProfiler^91^ package (v4.14.6), implementing the Fast GSEA (fgsea) algorithm with default parameters^92^. The Benjamini-Hochberg correction method was applied to adjust for multiple comparisons for differential expression and pathway analyses.

### 1-MDa *in vivo* microdialysis

Stereotaxic surgery and 1-MDa *in vivo* microdialysis was performed as previously described^25^. Mice were anesthetized with 2% isoflurane, and the head was shaved and fixed in a stereotaxic apparatus (Stoelting Co.). Skull position was leveled, and a hole was drilled at bregma −3.1 mm, 2.5 mm lateral to target the left caudal hippocampus. Another hole was made diagonal to the first to position an anchoring bone screw. After meninges were removed, an AtomosLM Guide Cannula (PEG-12, Eicom) was inserted at a 12° angle, 1.2 mm below the brain surface at the target location. The cannula was secured, and skin was closed using dental cement and surgical adhesive glue. An AtmosLM Dummy Cannula (PED-12, Eicom) was inserted into the guide cannula and secured with a plastic cap nut as the animal recovered in a clean cage on a heating pad. Approximately 12 hours following surgery, 1-MDa *in vivo* microdialysis probes (AtmosLM; Emicon) were inserted into caudal hippocampus. To calibrate the system prior to probe insertion, inlet tubing (FEP tubing 0.65 mm OD x 0.12 mm ID, BASi) was connected to a syringe pump (KdScientific) that perfused artificial cerebrospinal fluid (aCSF) containing, in mM: 1.3 CaCl_2_, 1.2 MgSO_4_, 3 KCl, 0.4 KH_2_PO_4_, 25 NaHCO_3_, 122 NaCl, pH 7.35, at a rate of 1.2 μl/min. The outlet tubing was connected to a peristaltic pump (MAB 20, SciPro), which was carefully calibrated to obtain a pull rate between 0.9-1.1 μl/min. Probe integrity was tested, and once passing initial quality control, it was connected to the inlet and outlet ports. The push-pull system was connected to the probe and allowed to equilibrate prior to implanting in mouse brain. The probe was secured with a cap-nut, and mice were placed in a Raturn (Stand-Alone Raturn System, BASi) and tethered by a loose collar to enable free movement while preventing tangling of microdialysis tubing. Dialysate from brain ISF was collected hourly at ∼1.0 μl/min into a refrigerated fraction collector (MAB 85 Fraction Collector, SciPro) and frozen at −80°C after collection for pooling and subsequent measurement. In microdialysis experiments involving LPS administration, mice were given a single i.p. injection of LPS (5 mg/kg in PBS), and samples were collected for an additional 10hrs. For all experiments, mice were given *ad libitum* access to food and water and were kept under constant light conditions to minimize the impact of circadian protein flux, as previously described^93^.

### Protein analysis from ISF dialysates

Hippocampal ISF dialysate samples were thawed on ice and pooled for measurements. For WT and TIMP2 KO comparisons, samples representing the stable 24-hour steady-state period were pooled within each mouse for subsequent measurements. For LPS experiments, the period representing a stable baseline over a 12-hour period were pooled within each mouse to serve as the pre-LPS sample per mouse. Samples corresponding to hours six to ten following LPS treatment were pooled within each mouse to serve as the post-LPS treatment samples per mouse. For all experiments, 230 μl of pooled sample was concentrated approximately 6-fold by centrifugation at 14,000 x g for 75 min at 4C with a concentrator unit (Amicon Ultra-0.5 Centrifugal Filter Units (MilliporeSigma). 25 μl of concentrated sample was loaded on a 96-well plate using the Olink Target Mouse Exploratory Panel, encompassing cytokines and chemokines (Olink, Uppsala Sweden) in coordination with the Human Immune Monitoring Center (ISMMS). Data was represented as Normalized Protein EXpression (NPX) unit on a log2 scale.

## Statistical analysis

Statistics were performed using GraphPad Prism version 10 (GraphPad Software) or R programming environment (version 4.4.2 or version 4.2.0) using tests described in figure legends. All relevant statistical tests were two-sided.

## Acknowledgements

We would like to thank Anna Podlesny-Drabiniok, Michael Sewell, Raphael Kubler, and Mikaela Rosen for advice related to snRNA-sequencing analysis, Eva Czirr for adult microglia isolation and culturing advice, Hanxiao Liu and Jeffrey Zhu for technical support, and Swastik P. G. for morphology advice. We thank Sanjana Shroff and Kristin Beaumont for assistance regarding snRNAseq (Single Cell Genomics Core, ISMMS), the Human Immune Monitoring Center (ISMMS) for Olink measurements, and Deanna Benson and Nikos Tzavaras of the Microscopy CoRE and Advanced Bioimaging Center (ISMMS) for advice related to brain phagocytosis imaging and analysis using Imaris software. Components of some figure panels used BioRender content (https://BioRender.com/7o21lrw). We acknowledge computational and data resources provided by Scientific Computing and Data (ISMMS), supported by the National Center for Advancing Translational Sciences, UL1TR004419, and NIH grants S10OD026880 and S10OD030463. This work was supported by the National Institute on Aging (R01AG061382 (JMC), RF1AG072300 (JMC),1F31AG079604-01A1 (BMH), T32AG049688 (BMH, SMP), and R01AG061382-02S1 (JMC, SMP)).

## Author contributions

BMH and JMC conceived and designed experiments, performed data analysis and interpretation, and wrote the manuscript. BMH performed experiments. JMC supervised the research. ACF generated tissue related to neuron-specific TIMP2 deletion, and some tissue for WT and TIMP2 KO mice, with percentage area and activation analysis for select cohorts, aided by AP. SMP dissected hippocampi for sequencing experiments and provided additional expertise to guide sequencing analysis. SFP performed morphology analysis and assisted with tissue isolation for adult microglial cultures. All authors provided input and approved the paper.

---

**Supplementary Figure 1.**
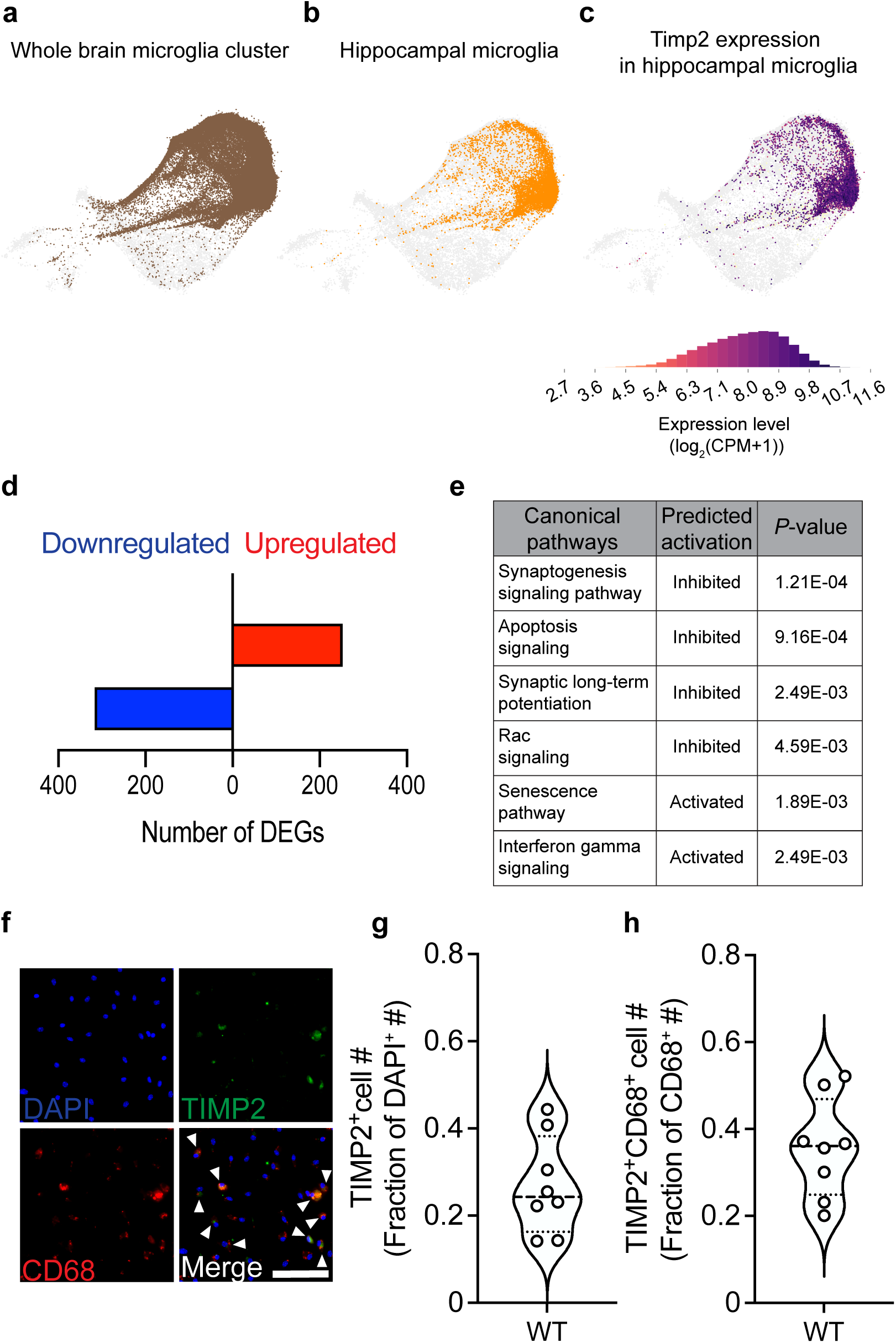
Microglia express TIMP2 protein near lysosomes. **(a)** Data from Allen Brain Cell scRNAseq and MERFISH Atlas representing whole brain microglia (brown), with subsets corresponding to **(b)** hippocampal microglia (orange), and **(c)** TIMP2 expression within hippocampal microglia. **(d)** Number of upregulated and downregulated DEGs in TIMP2 KO microglia relative to WT microglia (N = 7 samples per genotype, with each sample representing a pool of 5 pups (P6-P7); sex-matched). **(e)** IPA canonical pathways based on upregulated and downregulated DEGs in TIMP2 KO and WT microglia from panel (d), with corresponding predicted inhibition or activation with *P*-values. **(f)** Representative images from WT primary microglia cultures stained by ICC with DAPI, as well as anti-TIMP2 and anti-CD68 antibodies for overlap analyses (N = 8 mice, 5-6 month-old mice; sex-matched; 2 independent experiments combined; scale bar = 100 μm), with **(g)** quantification of the number of TIMP2^+^ cells as a fraction of DAPI^+^ cells, or **(h)** the number of TIMP2^+^CD68^+^ cells as a fraction of cells expressing CD68. Violin plots of median with quartiles.

**Supplementary Figure 2.**
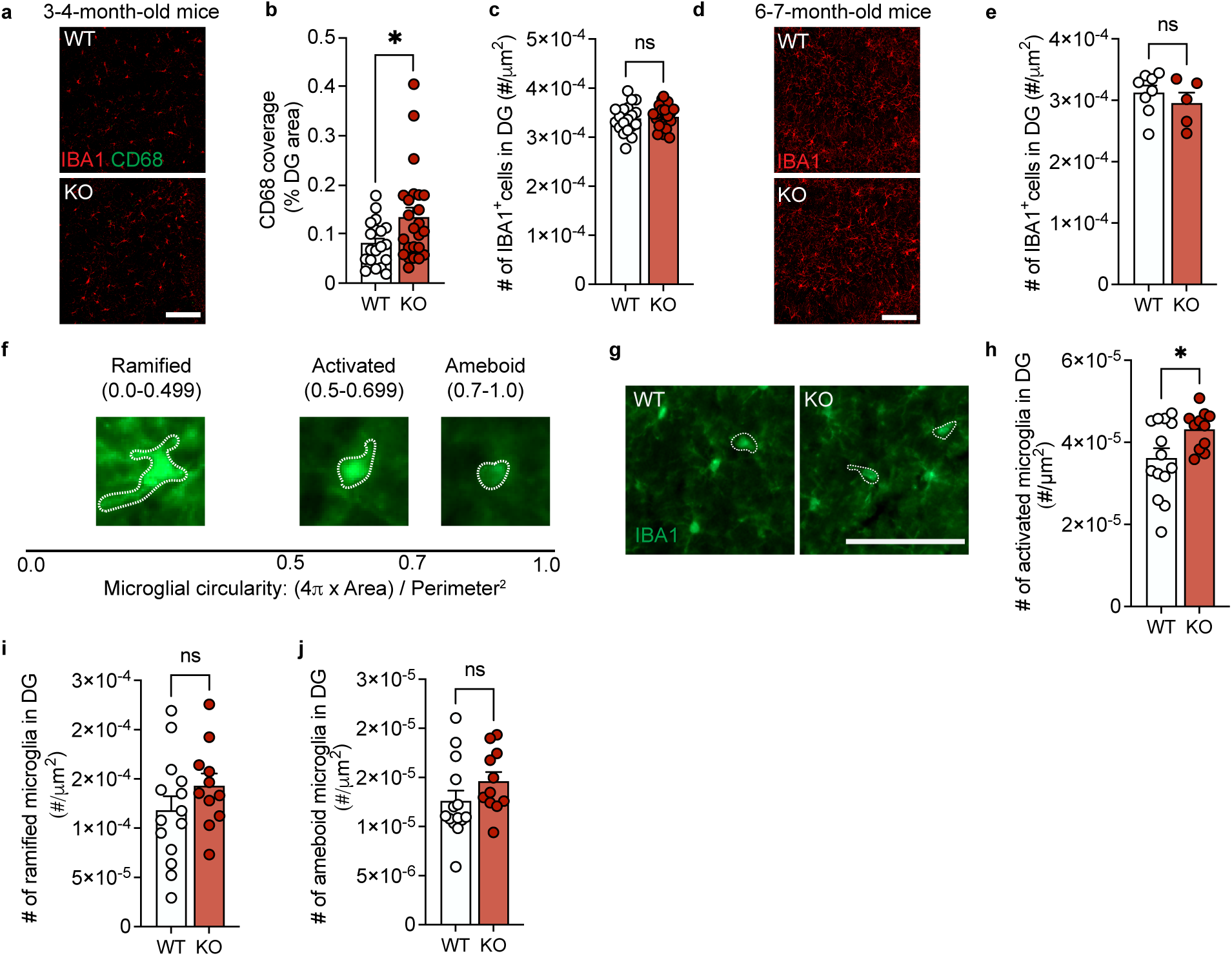
TIMP2 KO mice exhibit altered state and morphology associated with activation. **(a)** Representative confocal images of DG from 3-4-month-old WT and TIMP2 KO mice stained with anti-CD68 and anti-IBA1 antibodies (scale bar = 100 μm) with **(b)** quantification of thresholded DG area covered by CD68^+^ staining and **(c)** the number of IBA1^+^ cells normalized to DG area (N = 9-13 mice per group; sex-matched; mean ± SEM). **(d)** Representative confocal images of DG from 6-7-month-old WT and TIMP2 KO mice stained with anti-IBA1 antibody (scale bar = 100 μm) with **(e)** quantification of the number of IBA1^+^ cells normalized to DG area (N = 5-8 mice per group; female; mean ± SEM). **(f)** Representation of morphology analysis based on microglia circularity, with representative images for each morphological state according to circularity scale. **(g)** Representative microscopy images of DG from 6-7-month-old WT and TIMP2 KO mice stained with anti-IBA1 antibodies, with white dashed lines outlining microglia with “activated” morphology and corresponding (scale bar = 100 μm) **(h)** quantification of number of microglia with “activated” microglia (circularity = 0.5-0.699) per DG area. **(i)** Quantification of the number of microglia with “ramified” morphology (circularity = 0.0-0.499) per DG area, and **(j)** the number of microglia with “ameboid” microglia (circularity = 0.7-1.0) per DG area. (N = 11-14 mice per group; sex-matched; mean ± SEM). \**P* < 0.05; Student’s t test; n.s., not significant.

**Supplementary Figure 3.**
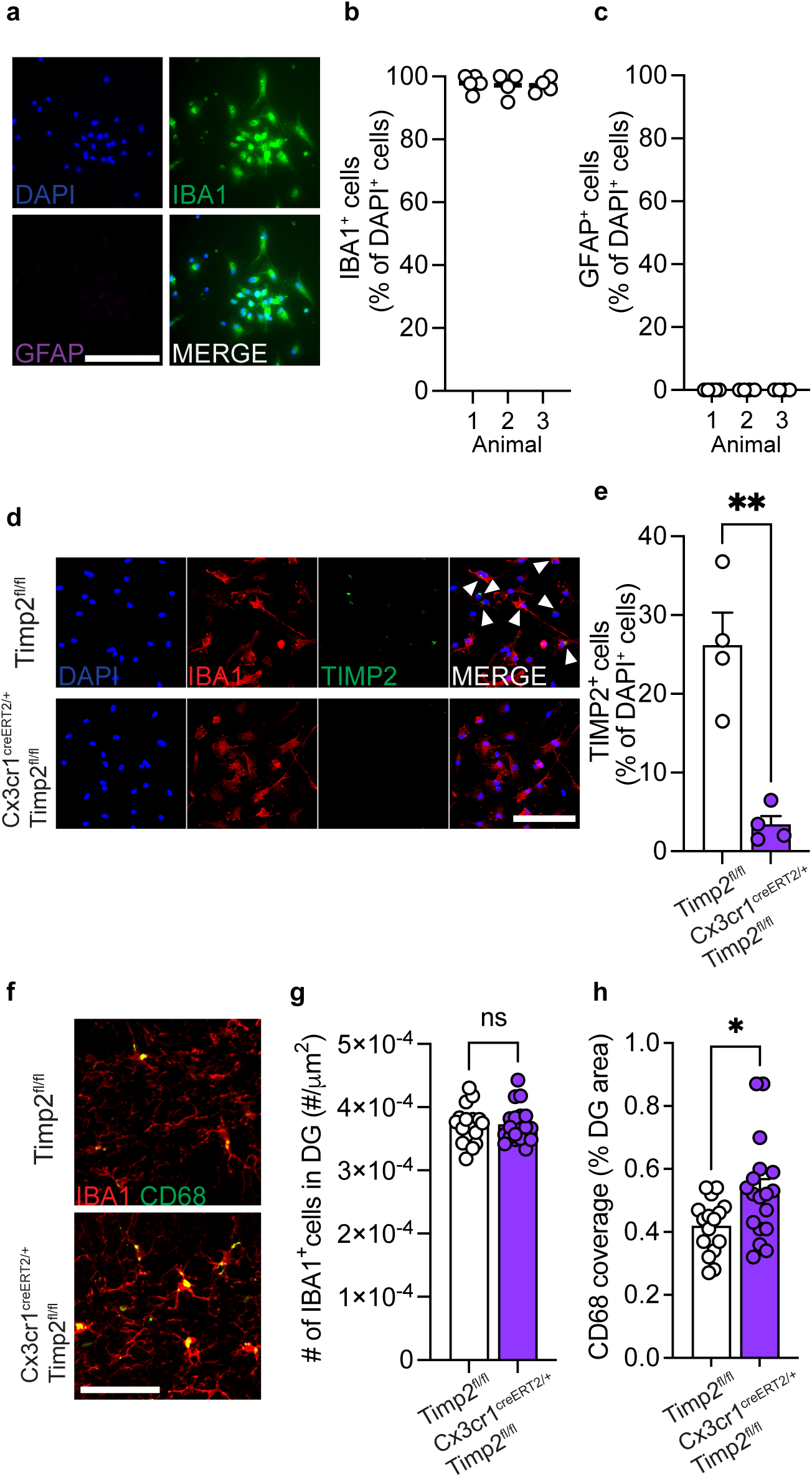
Purity of primary microglia cultures and efficiency of microglia *Timp2* deletion. **(a)** Representative microscopy images of primary microglia cultures from WT mice stained with DAPI, anti-IBA1, and anti-GFAP antibodies (scale bar = 100 μm) with **(b)** quantification of IBA1^+^ cells as a percentage of DAPI^+^ cells or **(c)** quantification of GFAP^+^ cells as a percentage of DAPI^+^ cells (N = 4-5 replicates per mouse from N = 3 mice). **(d)** Representative confocal images from primary microglia cultures from 5-month-old Timp2^fl/fl^ control and Cx3cr1^CreERT2/+^; Timp2^fl/fl^ littermates stained with DAPI, anti-IBA1 antibody, and anti-TIMP2 antibody (arrowheads indicate TIMP2^+^ microglia; N= 4 mice per group; sex-matched; scale bar = 100 μm) with **(e)** quantification of TIMP2^+^ cells as a percentage of DAPI^+^ cell number (mean ± SEM). **(f)** Representative confocal images of DG from 6-7-month-old TIMP2^fl/fl^ control and Cx3cr1^CreERT2/+^;Timp2^fl/fl^ littermates stained with anti-IBA1 and anti-CD68 antibodies (N= 17-19 mice per group; sex-matched; scale bar = 50 μm) with **(g)** quantification of the number of IBA1^+^ cells per DG area and **(h)** quantification of DG area covered by CD68^+^ staining (mean ± SEM). \**P* < 0.05; \*\**P* < 0.01; n.s., not significant. Student’s t test.

**Supplementary Figure 4.**
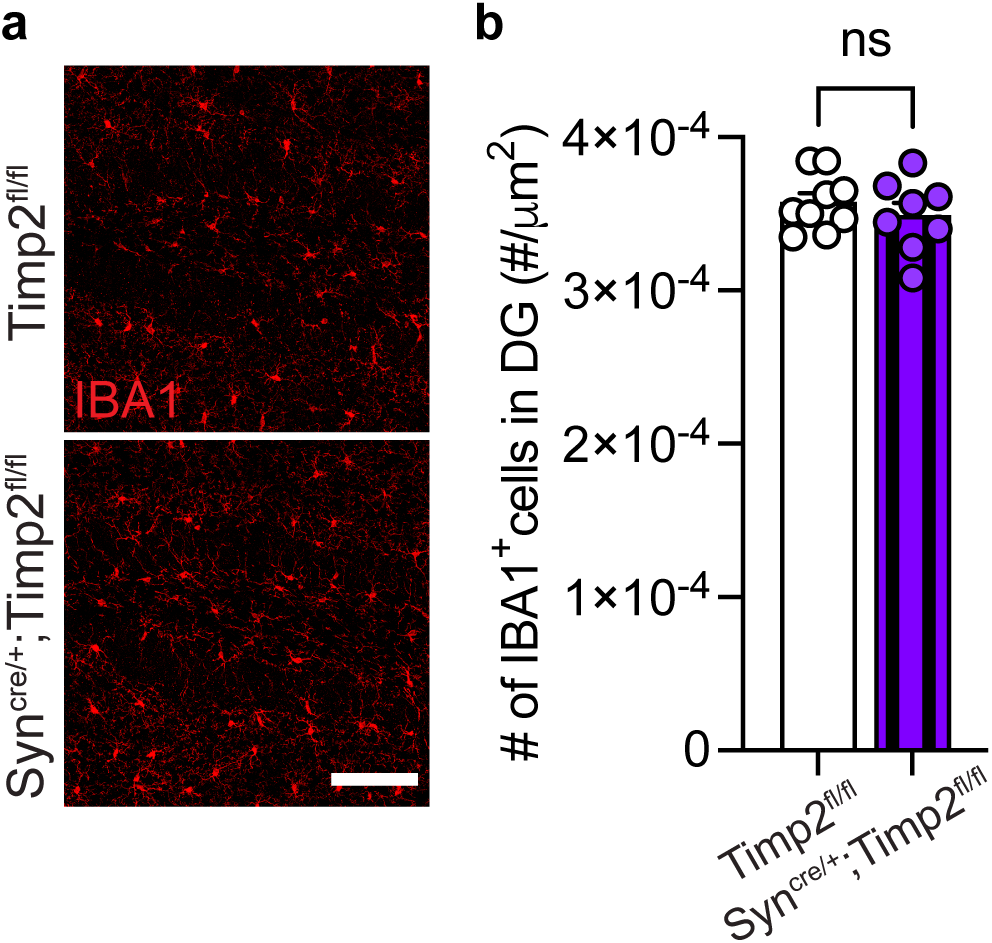
Neuronal deletion of TIMP2 does not alter number of microglia in the dentate gyrus. **(a)** Representative confocal images of DG from *Timp2*^fl/fl^ and Syn^Cre/+^;*Timp2*^fl/fl^ littermates stained with anti-IBA1 antibody with corresponding **(b)** quantification of the number of IBA1^+^ cells normalized to DG area (N = 8-9 mice per group, 2-3-month-old female mice; scale bar = 100 μm). mean ± SEM.; n.s., not significant. Student’s t test.

**Supplementary Figure 5.**
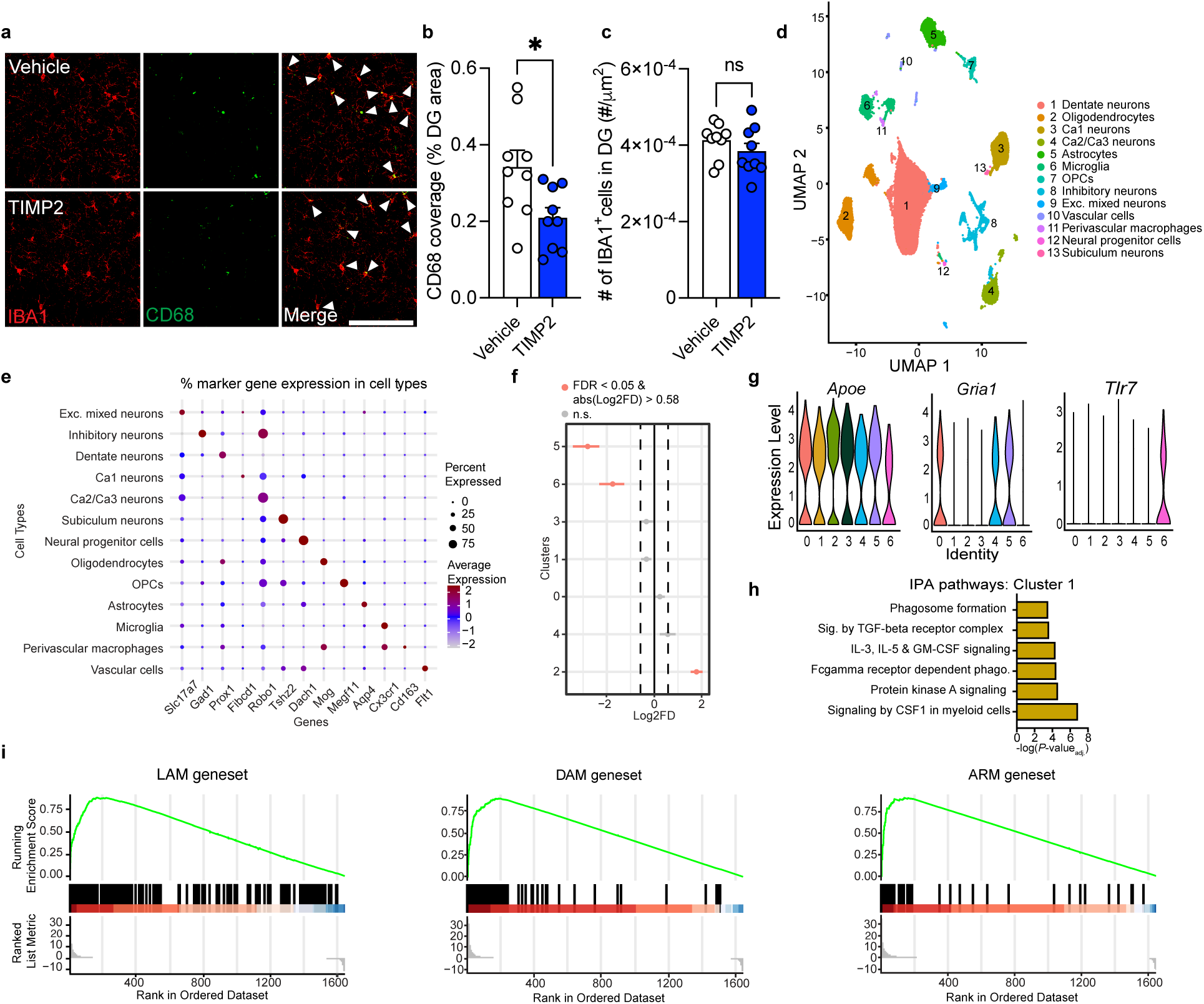
Systemic TIMP2 treatment in aged mice alters microglial state and shifts microglial subclusters. **(a)** Representative confocal images of DG from 20-month-old WT mice treated systemically with TIMP2 or vehicle, stained with anti-IBA1 and anti-CD68 antibodies (scale bar = 100 μm; white arrowheads indicate IBA1^+^CD68^+^ cells), with corresponding **(b)** quantification of the percentage of DG area covered by CD68^+^ staining and **(c)** quantification of the number of IBA1^+^ cells normalized to DG area (N = 9 male mice per group; mean ± SEM). **(d)** Uniform Manifold Approximation and Projection (UMAP) plot of hippocampal nuclei isolated from 20-month-old WT mice treated systemically with vehicle or TIMP2 (N = 4 male mice per group combined as 2 samples per group). **(e)** Dot plot of scaled expression of selected marker genes across annotated cell types. Dot size represents percentage of cells expressing the gene within each cell type, while color indicates the scaled average expression level of the gene across cells in the respective cluster. **(f)** Differences in cell proportions for each cluster between treatment conditions. Significant clusters are denoted in salmon (FDR < 0.05. Log_2_ fold-enrichment > 0.58, relative to vehicle-treated mice). **(g)** Violin plots of selected marker genes across microglial subclusters from Fig. 5h. **(h)** Significant canonical pathways from IPA for marker genes significantly upregulated in subcluster 1. **(i)** GSEA enrichment plots for LAM, DAM, and ARM gene sets based on the ranked list of microglial genes (TIMP2 vs. vehicle). Green lines represent running enrichment score, with vertical black bars indicating the position of genes from indicated gene sets in the ranked list and gene ranking direction indicated by color bar (red = upregulated, blue = downregulated).

